# Genetic dissection of crossover mutants defines discrete intermediates in mouse meiosis

**DOI:** 10.1101/2022.08.10.503530

**Authors:** Tolkappiyan Premkumar, Lakshmi Paniker, Rhea Kang, Mathilde Biot, Ericka Humphrey, Honorine Destain, Isabella Ferranti, Iyinyeoluwa Okulate, Holly Nguyen, Vindhya Kilaru, Melissa Frasca, Parijat Chakraborty, Francesca Cole

**Author notes:** Correspondence @ColeFrancesca.

## Abstract

Crossovers, the exchange of homolog arms, are required for accurate segregation during meiosis. Studies in yeast have established that the single end invasion intermediate is highly regulated to ensure crossover distribution. Single end invasions are thought to differentiate into double Holliday junctions that are resolved by MutLgamma (MLH1/3) into crossovers. Currently, we lack knowledge of early steps of mammalian crossover recombination or how intermediates are differentiated in any organism. Using comprehensive analysis of recombination and cytology, we infer that polymerized single-end invasion intermediates and nicked double Holliday junctions are crossover precursors in mouse spermatocytes. In marked contrast to yeast, MLH3 plays a structural role to differentiate single end invasions into double Holliday junctions with differentially polymerized 3’ ends. Therefore, we show independent genetic requirements for precursor formation and asymmetry with regard to 3’ end processing, providing mechanistic insight into crossover formation and patterning.

## Introduction

The fundamental mandate of meiosis is to generate haploid gametes from diploid precursors by first segregating parental chromosomes (homologs) and then sister chromatids. Homolog segregation is a challenge because chromosomes must be physically connected for accurate segregation, but only identical sister chromatids are tethered by cohesion. The solution in most organisms, including humans and mice, is to ensure homologous recombination as a crossover (CO) occurs between each homolog pair (Gray and Cohen, 2016; Hunter, 2015) while maintaining sister chromatid cohesion to physically tether homologs. Failure to ensure crossing over causes aneuploidy, and in humans, even mild CO defects cause infertility, miscarriages, and constitutional aneuploidies such as Down syndrome (Hassold and Hunt, 2001).

Meiotic recombination is initiated by SPO11-induced DNA double-strand breaks (DSBs) that form preferentially at hotspots (Keeney et al., 2014). DSBs undergo resection that requires 5’è3’ exonucleolytic digestion by Exo1 in yeast, but not in mouse (Mimitou and Keeney, 2018; Yamada et al., 2020; Zakharyevich et al., 2010). Resected 3’ overhangs form nucleoprotein filaments with DMC1 and RAD51 that engage in strand invasion to form a nascent three stranded D-loop (Gray and Cohen, 2016) (**Figure 1A**). COs are designated in early meiotic prophase (Borner et al., 2004), a step that in yeast requires Zip3 and MutSgamma, among other proteins and can be physically detected as a metastable strand exchange intermediate, the single-end invasion (SEI) (Hunter and Kleckner, 2001). These SEIs are thought to undergo second end capture to become double Holliday junctions (dHJs) that are ultimately resolved as COs (Hunter, 2015). In mammals, the Zip3 ortholog RNF212 stabilizes MutSgamma at recombination sites and is thought to be similarly required for CO designation (Qiao et al., 2014; Rao et al., 2017; Reynolds et al., 2013). HEI10, an E3 ubiquitin ligase antagonizes RNF212’s role in MutSgamma stabilization, but is also required for CO maturation, the step in which dHJs are resolved to form COs (Qiao et al., 2014; Ward et al., 2007). Proper CO maturation requires MutLgamma, a heterodimer MLH1 and MLH3 (Gray and Cohen, 2016), which is thought to function as a dHJ resolvase. Congruent with this model, MLH1 and MLH3 accumulate as foci at CO sites in mammals and MutLgamma-derived COs require the nuclease function of MLH3 (Gray and Cohen, 2016; Nishant et al., 2008; Raghavan, 2019; Toledo et al., 2019). Indeed, reconstituted MutLgamma complex containing a nuclease-dead allele of MLH3 lacks nicking activity in vitro (Cannavo et al., 2020; Kulkarni et al., 2020). When MutLgamma is absent, only 1 – 2 COs are formed per mouse spermatocyte and can be detected as connected homologs (bivalents) in metaphase (Guiraldelli et al., 2013; Lipkin et al., 2002; Ward et al., 2007; Zelazowski et al., 2017). These COs are likely produced by structure selective endonucleases (SSNs) such as MUS81 (Holloway et al., 2008).

**Figure 1.**
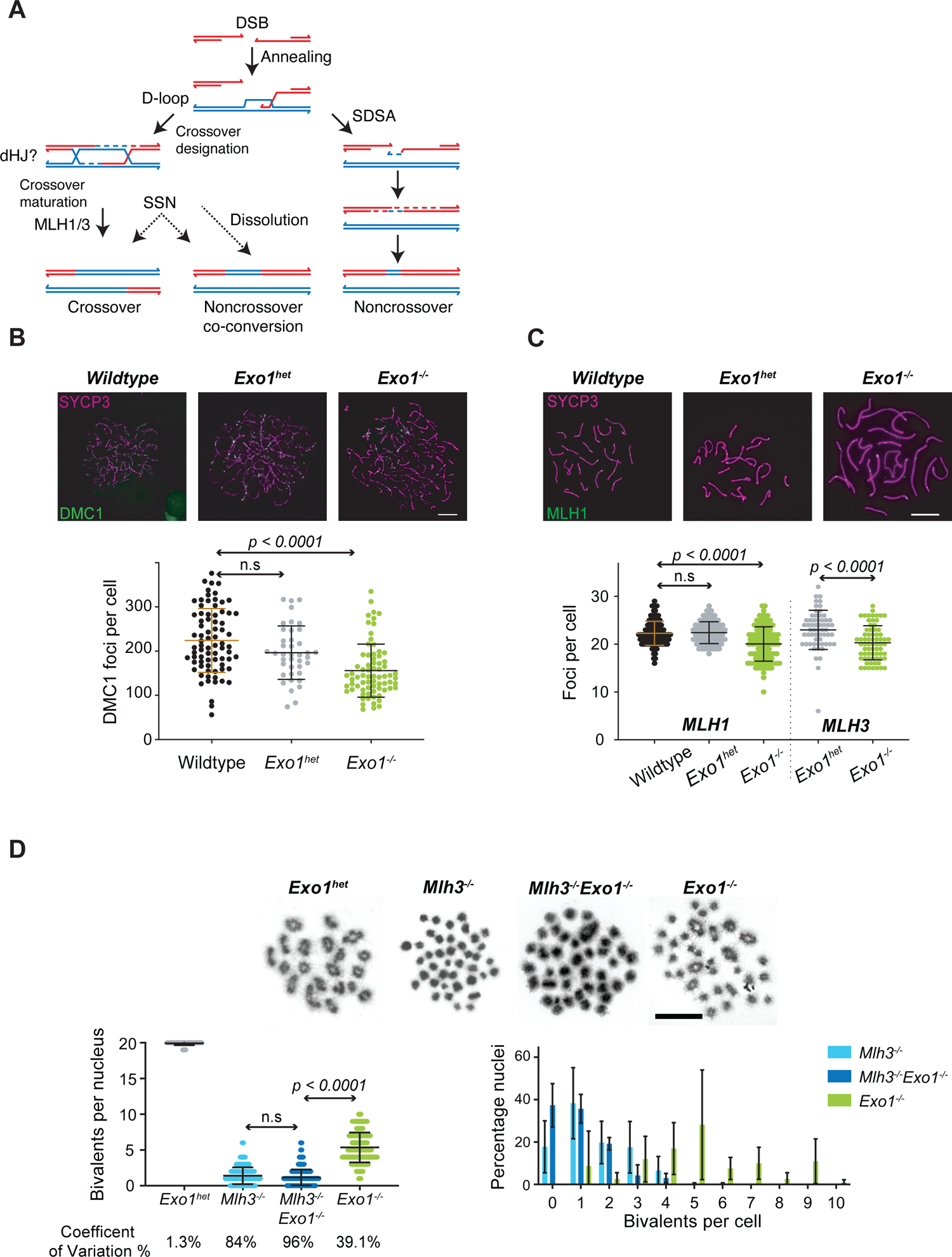
*Exo1^-/-^* spermatocytes have reduced recombination intermediates and fewer bivalents most of which require MLH3. **(A)** Current model of mammalian DSB repair in meiosis. The double Holliday junction (dHJ) is inferred based upon work in budding yeast. SDSA, synthesis-dependent strand annealing; SSN, structure-selective endonuclease. **(B)** Representative immunofluorescence images of zygotene stage spermatocytes of the indicated genotypes stained for SYCP3 (magenta) and DMC1 (white). Scatter plot of DMC1 foci per cell (y-axis) with error bars of the mean ± SD in WT (N=3, 224 ± 72.2), *Exo1^het^* (N=2, 196.5 ± 60.45), and *Exo1^-/-^* (N=4, 155.8 ± 60). P-values were determined by Kruskal-Wallis test with Dunn’s multiple comparison correction. *Exo1^-/-^* spermatocytes showed fewer DMC1 foci per cell on average in zygonema as compared to WT and *Exo1^het^* spermatocytes. **(C)** Representative immunofluorescence images of pachytene stage spermatocytes of the indicated genotypes stained for SYCP3 (magenta) and MLH1 (white). Scatter plot of MLH1 and MLH3 foci per cell as above in WT (MLH1, N=3, 22.3 ± 2.5), *Exo1^het^* (MLH1, N=3, 22.4 ± 2.3), *Exo1^-/-^*(MLH1, N=3, 20.0 ± 3.6) and *Exo1^het^* (MLH3, N=2, 22.9 ± 4.1), *Exo1^-/-^* (Mlh3, N=3, 20.3 ± 3.6). P-values were determined by Kruskal-Wallis test with Dunn’s multiple comparison correction (MLH1) and Mann-Whitney, two-tailed (MLH3). *Exo1^-/-^* spermatocytes showed fewer MLH1 and MLH3 foci per cell on average in pachynema as compared to WT and *Exo1^het^* spermatocytes. **(D)** Representative metaphase images of spermatocytes from the indicated genotypes stained with Giemsa. Left, scatter plot of bivalent counts per cell (y-axis) with error bars of the mean ± SD in *Exo1^het^* (N=3, 19.9 ± 0.3), *Mlh3^-/-^* (N=5, 1.4 ± 1.2), *Exo1^-/-^Mlh3^-/-^*(N=4, 1.2 ± 1.1), and *Exo1^-/-^* (N=4, 5.3 ± 2.0). P-values were determined by Kruskal-Wallis test with Dunn’s multiple comparison correction. Right, histogram of the distribution of bivalents per cell of the indicated genotypes. The x-axis shows the number of bivalents per cell and the y-axis shows the average percentage of total nuclei with the indicated number of bivalents. The two graphs also visually depict the Coefficient of Variation of bivalents per genotype *Mlh3^-/-^* (84%), *Exo1^-/-^Mlh3^-/-^* (96%), *Exo1^-/-^* (39.1%), and *Exo1^het^* (1.3%). *Exo1^-/-^* spermatocytes had fewer bivalents than *Exo1^het^* but more than *Mlh3^-/-^* spermatocytes. *Exo1^-/-^Mlh3^-/-^* analysis shows the majority of the residual bivalents in *Exo1^-/-^*require MLH3.

Most DSBs in mammals (∼90%) are repaired as noncrossovers (NCOs) - short patch-like repair products that do not involve exchange of homolog arms (Cole et al., 2012b). Most NCOs form via synthesis-dependent strand-annealing (SDSA) (Allers and Lichten, 2001) a process that does not involve formation of a dHJ. In wildtype (WT) mouse and human spermatocytes, NCOs mostly incorporate only one polymorphism from the donor, hence referred to as singletons (Cole et al., 2010; Jeffreys and May, 2004). In mammals, only ∼10% of DSBs are thought to be designated to form COs during early meiotic prophase. In the absence of MLH3, these designated COs are acted upon by alternative repair pathways to produce long NCOs called co-conversions that involve multiple polymorphisms (Zelazowski et al., 2017).

In yeast, Exo1 plays an indispensable and nuclease-independent role in CO maturation by MutLgamma (Gioia et al., 2021; Keelagher et al., 2011; Zakharyevich et al., 2010). Consonant with this function, EXO1 can stimulate MutLgamma nicking activity in association with MutSgamma and RFC-PCNA in vitro (Cannavo et al., 2020; Kulkarni et al., 2020). *Exo1^-/-^* mice are sterile and have ∼80% fewer COs, but only ∼10% fewer MLH1/3 foci than WT spermatocytes (Kan et al., 2008; Wei et al., 2003). Mice harboring the nuclease-dead allele of *Mlh3* (*Mlh3^DN^*) also show disproportionately fewer COs despite proficient MLH1/MLH3 focus formation (Toledo et al., 2019). The CO promoting role of EXO1 is also nuclease-independent as mice carrying a nuclease dead allele of *Exo1* (*Exo1^D173A^*) are fertile and proficient in CO formation (Wang et al., 2022). Unlike yeast, the CO phenotype in mice lacking EXO1 or MutLgamma are not similar raising the question of whether mammalian EXO1 and MLH3 play distinct roles in CO intermediate formation.

Much of our understanding of CO mechanisms is extrapolated from studies in yeast. This is in part due to detection of early CO precursor intermediates (Hunter and Kleckner, 2001; Schwacha and Kleckner, 1995) and retention of extensive heteroduplex when mismatch repair is compromised by deletion of Msh2 (Ahuja et al., 2021; Marsolier-Kergoat et al., 2018; Martini et al., 2011). However, CO precursors haven’t been isolated in mammals and there are many unique features of recombination including infrequently retained heteroduplex (Peterson et al., 2020) As such, we lack understanding of the early steps of CO recombination. Using fine-scale, comprehensive analysis of recombination outcomes and cytology of 12 different mutant or mutant combinations, we provide evidence for two transient states of designated CO intermediates and define the roles of EXO1 and MLH3 in mouse spermatocytes. Key findings are that 1) the designated CO intermediate polymerizes from a single invaded 3’ end likely forming an extended SEI intermediate; 2) MLH3 plays an unexpected structural role to differentiate the SEI intermediate into a dHJ; 3) CO precursor processing is asymmetric with regard to extent of 3’ polymerization, which can explain CO biased resolution and patterning; and 4) the MLH3-structure dependent dHJs are more likely to be resolved as COs by SSNs.

## Results and Discussion

### Mouse spermatocytes lacking EXO1 have disproportionately fewer COs than recombination intermediates

In budding yeast, loss of MutLgamma and Exo1 show similar phenotypes (Zakharyevich et al., 2010) precluding epistatic analysis of null alleles. In mice, however, inactivation of *Exo1* does not share a similar phenotype to *Mlh1^-/-^* or *Mlh3^-/-^* (Kan et al., 2008). We find that *Exo1^-/-^* spermatocytes have fewer DMC1 foci in zygonema (155.8 ± 60) than WT or *Exo1^het^* (224.0 ± 72.2 and 196.5 ± 60.45, respectively) (**Figure 1B**). By contrast, loss of MLH3, shows equal numbers of recombination intermediates (RAD51 and gammaH2AX) in early zygonema as WT, although there are fewer RAD51 foci in late zygonema in *Mlh3^-/-^* spermatocytes (Toledo et al., 2019). Similarly, there are lower RPA2 foci in *Mlh3^-/-^* spermatocytes in zygonema, but equivalent numbers in pachynema (Zelazowski et al., 2017). Lower numbers of DMC1 foci could reflect reduced overall DSBs, reduced resection, or an overall reduction in lifespan of the DMC1 nucleoprotein filament. Exo1 is required for most meiotic DSB resection in budding yeast (Garcia et al., 2011; Keelagher et al., 2011; Mimitou and Keeney, 2018; Zakharyevich et al., 2010), however loss of EXO1 nuclease activity lowers DSB resection by only 10% in the mouse (Paiano et al., 2020; Yamada et al., 2020). While it is possible that *Exo1^-/-^* spermatocytes have fewer DSBs, this is not found in budding yeast (Keelagher et al., 2011; Zakharyevich et al., 2010). Further, axis lengths, which are usually proportional to meiotic DSB frequency (Keeney et al., 2014) are unchanged between WT and *Exo1^-/-^* pachytene spermatocytes (**Figure S1A**). Finally, total recombination frequency measured at the *59.5* hotspot in *Exo1^-/-^* diplotene spermatocytes (196.7 per 10,000 haploid genome equivalents) is similar to both WT (213.3), *Mlh3^-/-^* (194.6), and several other mutants that do not show alteration in meiotic DSB frequency (**Table 1**, **Figure S1B**) (Rao et al., 2017; Ward et al., 2007). We suggest reduced DMC1 foci in *Exo1^-/-^*spermatocytes reflects a combination of mildly reduced resection and lifespan of the DMC1 nucleoprotein filament.

The 26% fewer DMC1 foci in *Exo1^-/-^* spermatocytes should be easily mitigated by CO homeostasis (Martini et al., 2006), which in mouse spermatocytes ensures CO number despite mild fluctuations in the overall number of early recombination intermediates (Cole et al., 2012a). However similar to earlier reports in oocytes (Kan et al., 2008), we find only 20.0 ± 3.6 MLH1 foci per cell on average in *Exo1^-/-^*spermatocytes as compared to 22.3 ± 2.5 and 22.4 ± 2.3 in WT and *Exo1^het^*, respectively (**Figure 1C**). We find a similar ∼10% reduction in MLH3 foci. The lower number of MutLgamma foci could reflect a role(s) for EXO1 in CO homeostasis and in support of this model, *Exo1^-/-^*yeast have reduced CO interference (Gioia et al., 2021) – the observation that COs are further apart than expected by chance. Reduced CO interference is seen when CO homeostasis is compromised (Wang et al., 2015). However, we find that CO interference as measured by the coefficient of coincidence of MLH1 foci is unchanged in *Exo1^-/-^* spermatocytes as compared to wild type (**Figure S1C)**. Alternatively, EXO1 could play a role in the lifespan or stability of the MutLgamma complex, in support of this model, EXO1 physically associates with MLH1 (Tran et al., 2001), promotes MutLgamma nuclease activity in vitro (Cannavo et al., 2020; Kulkarni et al., 2020), and can increase the stability of MLH1 containing mismatch repair complexes (Amin et al., 2001). Given that CO interference is unchanged in *Exo1^-/-^* spermatocytes, we favor the latter model that lower MLH1 and MLH3 foci reflect reduced stability of the MutLgamma complex in the absence of EXO1.

Although there are substantial MLH1/MLH3 foci in *Exo1^-/-^* spermatocytes, most of these sites do not result in COs (Kan et al., 2008; Wei et al., 2003). We observed fewer bivalents per cell on average in *Exo1^-/-^*(5.3 ± 2.0) as compared to *Exo1^het^* spermatocytes (19.9 ± 0.3) (**Figure 1D**). Consistent with earlier reports, *Exo1^-/-^* showed a less severe reduction in bivalents than *Mlh3^-/-^* spermatocytes (Kan et al., 2008). This finding suggests that MLH3, and likely MutLgamma may be required for the majority of COs in *Exo1^-/-^* spermatocytes. Consistent with this hypothesis, *Mlh3* is epistatic to *Exo1* as *Exo1^-/-^Mlh3^-/-^* spermatocytes have similar number and distribution of bivalents per cell as that of *Mlh3^-/-^* spermatocytes (**Figure 1D**).

Variability in the number of bivalents per cell is low in spermatocytes with efficient CO maturation likely reflecting strong CO assurance. For example, in WT and *Exo1^het^* spermatocytes we found Coefficents of Variation (CoVs, a ratio of the standard deviation by the mean) of 2.7% and 1.3%, respectively. By contrast in *Mlh3^-/-^* and *Exo1^-/-^Mlh3^-/-^* spermatocytes the CoVs were 84% and 96%, respectively. The high cell to cell variation in the number of bivalents could reflect that SSNs do not generate COs from intermediates that are subject to CO assurance mechanisms. However, we found that 2/3 of COs in *Mlh3^-/-^* spermatocytes require RNF212, suggesting that they largely derive from CO designated and assured intermediates ((Reynolds et al., 2013) and R.K, F.C., unpublished data). Instead, the high variability of bivalents per cell likely reflects variability in the number of intermediates or in the number of intermediates that can be acted upon by SSNs. We favor the latter model as yeast lacking Mlh3 have wild-type levels of joint molecules (Arter et al., 2018; De Muyt et al., 2012; Zakharyevich et al., 2012), overall recombination activity is similar between WT, *Mlh3^-/-^*, and *Exo1^-/-^* spermatocytes (**Figure S1B**), and because CO interference is unchanged in *Exo1^-/-^* spermatocytes (**Figure S1C**). Given this model, it is interesting that we find an intermediate CoV for bivalents per cell in *Exo1^-/-^* spermatocytes of 39.1% (**Figure 1D**), suggesting a large fraction of these COs may derive from SSNs contradicting the simple hypothesis that most COs in *Exo1^-/-^* spermatocytes are MutLgamma-derived.

### *Exo1^-/-^* COs are dependent upon MLH3 and independent of EXO1 nuclease activity

To definitively determine the roles of EXO1 in CO formation, we compared recombination outcomes between mutants at the *59.5* hotspot on Chr19. Spermatocytes lacking COs apoptose at metaphase I causing azoospermia and infertility. To directly compare mutants with WT, we isolated spermatocytes at the end of meiotic prophase I (Late 4C). In WT sperm, we find that 67% of recombinants at *59.5* are COs (Zelazowski et al., 2017) (**Table 1, Figure S1B**). The remaining recombinants are singletons, NCO gene conversions that include only one polymorphism (29%) and co-conversions, NCO gene conversions that include two or more polymorphisms (4%). In support of our experimental approach, recombination outcomes are equivalent between sperm and isolated diplotene spermatocytes in WT (**Figure S1B**).

To determine the contribution of EXO1-dependent nuclease activity to crossing over, we compared recombination between *Exo1^-/-^* spermatocytes and a nuclease-deficient allele (*Exo1^nd^*) that replaces one of the five metal binding residues (D173A) and reduces nuclease activity by at least 98% in vitro (Myler et al., 2016; Shao et al., 2014; Sokolsky and Alani, 2000; Tran et al., 2002; Wang et al., 2022; Zhao et al., 2018). *Exo1^het^* and *Exo1^nd/nd^* spermatocytes showed 28% lower CO frequencies than WT spermatocytes at *59.5* (**Figure 2A**). However both *Exo1^het^* and *Exo1^nd/nd^* mice are fertile and global bivalents in metaphase are similar between *Exo1^het^* and WT spermatocytes (Wang et al., 2022) (**Figure 1D**). It is possible that in-depth analysis of recombination at *59.5* may reveal subtle haploinsufficient phenotypes for EXO1. To increase rigor, we limited any comparisons of *Exo1^-/-^* spermatocytes to *Exo1^het^*. *Exo1^-/-^*spermatocytes received 77.9% fewer COs compared to *Exo1^het^* (**Figure 2A, Table 1**), similar to the 73.4% fewer COs observed globally (**Figure 1D**) and supporting the concept that in depth analysis of *59.5* can reflect general recombination mechanisms. As expected from previous reports (Keelagher et al., 2011; Zakharyevich et al., 2010), this CO defect was not seen in *Exo1^nd/nd^* mice (Wang et al., 2022) (**Figure 2A, Table 1**) and they are fertile (Zhao et al., 2018). Taken together, these findings suggest the CO role of EXO1 in mice is nuclease-independent globally and at *59.5*.

**Figure 2.**
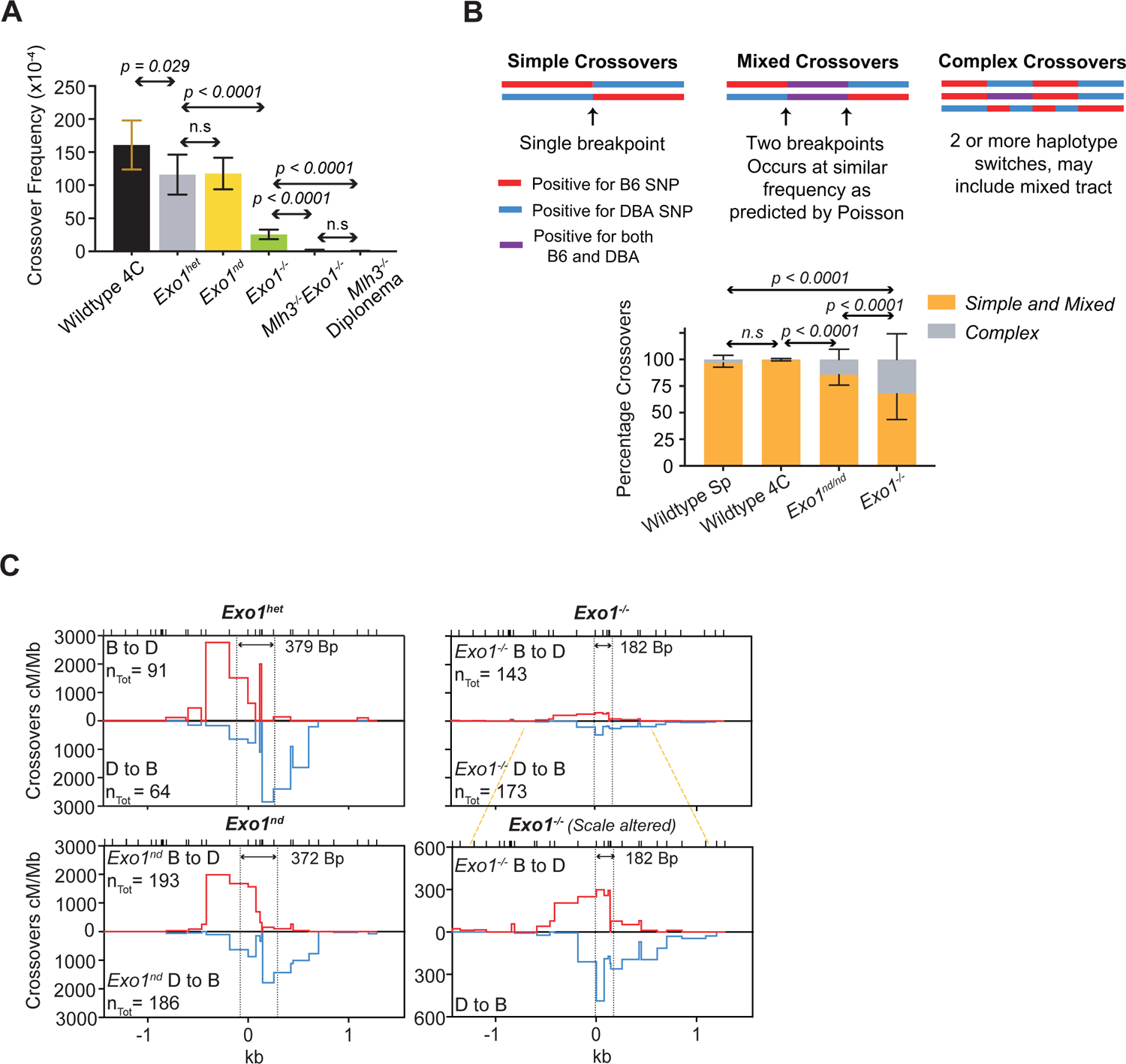
Meiotic COs are independent of EXO1 nuclease activity and residual COs in *Exo1^-/-^* spermatocytes have altered characteristics. **(A)** Poisson-corrected frequencies of COs at the *59.5* hotspot are plotted per 10,000 haploid genome equivalents (y-axis) with error bars of the mean ± SD in WT (N=4, 160.8 ± 37.1), *Exo1^het^* (N=4, 116.0 ± 30.2), *Exo1^nd/nd^* (N=4, 117.6 ± 23.9), *Exo1^-/-^* (N=6, 25.59 ± 7.32), *Mlh3^-/-^ Exo1^-/-^* (N=4, 0.77 ± 1.56), and *Mlh3^-/-^* (N=3, 0.28 ± 0.50). P-values were determined by Fisher’s exact test, two-tailed. *Exo1^nd/nd^*showed similar CO frequency as *Exo1^het^* spermatocytes, and both are fertile suggesting most COs are independent of EXO1 nuclease-activity. *Exo1^-/-^* had more residual COs compared to *Mlh3^-/-^* spermatocytes, and they require MLH3. **(B)** COs with one haplotype switch between parental genotypes were designated as simple COs. COs with one contiguous tract that scores positive for both parental genotypes were designated mixed COs. Mixed COs often arise when more than one simple CO is present in a sample. Mixed COs were found at the frequency expected for multiple COs per sample and were therefore included with simple COs for this analysis. COs with more than two haplotype switches that may involve tracts that are positive for both parental genotypes were designated as complex COs. Complex COs are likely derived from a single recombination event that experiences template switching or branch migration during CO formation and may include retained heteroduplex. Histogram of the percent of simple (yellow) and complex (gray) CO recombination patterns (y-axis) of the indicated genotypes. Frequency of complex COs in WT sperm (N=6, 2.5 ± 3.5), WT Late 4C spermatocytes (N=4, 0.4 ± 0.9), *Exo1^nd/nd^* (N=4, 14.4 ± 9.7), and *Exo1^-/-^* (N=6, 32.4 ± 24.1). P-values were determined by Fisher’s exact test, two-tailed. *Exo1^-/-^* spermatocytes had significantly more complex COs than WT and twice as many as *Exo1^nd/nd^*. **(C)** CO breakpoint frequency in centiMorgans per Megabase (cM/Mb) is plotted on the y-axis and CO breakpoint distribution plotted on the x-axis with the hotspot center set at 0 kb. The area under the curve represents total CO frequency. Note the CO breakpoint map for B to D COs (red, top) and D to B COs (blue, bottom) cluster on opposite sides of the hotspot center indicating reciprocal CO asymmetry due to most DSBs forming on the B chromosome. Vertical dotted lines mark the average location of CO breakpoints by cumulative distribution with the difference between them estimating the mean CO-dependent gene conversion tract length. *Exo1^-/-^*spermatocytes have a high frequency of CO breakpoints near the hotspot center and concomitantly shorter estimated gene conversion tract length compared to *Exo1^het^*or *Exo1^nd/nd^* spermatocytes. Top ticks, genotyped polymorphisms. n_Tot_, total number of COs mapped.

*Exo1^-/-^* had 91-fold higher COs at *59.5* compared to *Mlh3^-/-^* spermatocytes (**Figure 2A**). Similar to the genetic dependence for global crossing over, *Mlh3* was epistatic to *Exo1* for COs at *59.5* as *Exo1^-/-^Mlh3^-/-^* spermatocytes had similar CO frequency to *Mlh3^-/-^* (**Figure 2A, Table 1**). Taken together, COs in *Exo1^-/-^* spermatocytes require MLH3 and are independent of EXO1 nuclease activity.

### COs in *Exo1^-/-^* spermatocytes are more complex and show altered breakpoints

Most COs observed in WT and mutant spermatocytes show a single exchange point between parental haplotypes (simple COs) (**Figure 2B)**. However, we occasionally see COs that involve a contiguous tract that scores positive for both parental haplotypes. We refer to these as ‘mixed COs’ as the simplest interpretation is that these samples contain two independent simple COs that have different exchange points. In support of this interpretation, the number of mixed COs was not significantly different from the number of samples predicted to have multiple COs by Poisson approximation and the frequency of mixed COs is lower when fewer input molecules are used. We can observe complex COs with more than two haplotype switches that may involve tracts that are positive for both parental haplotypes. These complex COs cannot be explained as multiple events, but are likely caused by template switches or branch migration during CO formation and/or mismatch repair. These events are rare in WT spermatocytes or sperm (0.4 ± 0.9% and 2.5 ± 3.5%, respectively), however complex COs were more frequently seen in spermatocytes lacking EXO1 (32.4 ± 24.1%) (**Figure 2B)**. EXO1-dependent mismatch correction predominantly requires EXO1’s nuclease function (Goellner et al., 2015; Wang et al., 2022). In order to determine the contribution to complex COs from defective mismatch repair, we compared *Exo1^-/-^* and *Exo1^nd/nd^* spermatocytes (**Figure 2B**). We observed fewer complex COs in *Exo1^nd/nd^* spermatocytes (14.4 ± 9.7%). Together, the higher frequency of complex COs seen in *Exo1^-/-^* spermatocytes likely reflects both nuclease-dependent and -independent roles of EXO1, implying that template switching/branch migration is likely more frequent in *Exo1^-/-^* COs than that of *Exo1^nd/nd^* and WT.

Most meiotic DSBs at the *59.5* hotspot in a BxD F1 hybrid occur on the B chromosome. As such, most of the CO-dependent gene conversions alter the recipient B chromosome in favor of the donor D chromosome. As a consequence CO breakpoint distributions show reciprocal CO asymmetry (Jeffreys and Neumann, 2002; Zelazowski et al., 2017) (**Figure 2C**). The majority of B (recipient) to D (donor) CO breakpoints cluster to the left of the hotspot center (Top, red) and the majority of D to B breakpoints cluster to the right (Bottom, blue). Additionally, the distance between the midlines of the two CO breakpoint distributions reflects the average CO-dependent gene conversion tract. The estimated gene conversion tract length observed in *Exo1^het^* and *Exo1^nd/nd^* spermatocytes was ∼400bp similar to that of WT (**Figure 2C**) (Zelazowski et al., 2017). However, we observed that *Exo1^-/-^* spermatocytes had a lower estimated gene conversion tract length of 182bp. As *Exo1^nd/nd^* spermatocytes have similar estimated gene conversion tract length as *Exo1^het^*, it is unlikely that defective mismatch repair in *Exo1^-/-^* contributes to this phenotype.

This decrease in estimated gene conversion tract length in *Exo1^-/-^* spermatocytes may be caused by a decrease in the proportion of DSBs or CO designations on the B vs D homolog. However, we see 2.3- and 2.0-fold more NCOs on the B versus D chromosome in *Exo1^-/-^* and *Exo1^het^* spermatocytes, respectively, consistent with similarly high DSBs on B in both genotypes, and we still see reciprocal CO asymmetry in *Exo1^-/-^*spermatocytes despite the shorter apparent gene conversion tract. It is also possible that *Exo1^-/-^* spermatocytes have shorter CO-dependent gene conversions, however this should correlate with the length of all recombination outcomes derived from the designated CO intermediate and we do not see this (described below). Instead we favor a model that there is non-stereotypical positioning of CO breakpoints in *Exo1^-/-^* spermatocytes because we observe many CO breakpoints accumulating towards the center of the hotspot, where we rarely see breakpoints in *Exo1^het^* and *Exo1^nd/nd^* spermatocytes (p = 0.0005 and 0.0416, respectively Fishers Exact test, two-tailed, **Figure 2C**). The enrichment of CO breakpoints in the center of the hotspot may reflect altered branch migration of the double Holliday junction intermediate in *Exo1^-/-^* spermatocytes. In support of this model, the inferred positions of Holliday junctions are altered in budding yeast in the absence of MutLgamma or Exo1 (Marsolier-Kergoat et al., 2018). Alternatively the enrichment may be driven by differences in the endonucleolytic behavior of CO resolving enzymes.

### Most COs in *Exo1^-/-^* spermatocytes are derived from SSNs

*Exo1^-/-^* spermatocytes show higher CO levels than *Mlh3^-/-^*, both locally at *59.5* and globally in metaphase bivalents, and all these COs require MLH3 (**Figure 1D, 2A**), suggesting they derive from residual MutLgamma cleavage. Counterintuitively, the CoV of bivalent numbers per cell in *Exo1^-/-^*spermatocytes support a model that COs in *Exo1^-/-^* may arise via enzymatic activity of SSNs. A cohesive interpretation of these disparate findings could be that the residual COs in *Exo1^-/-^* spermatocytes require MutLgamma structurally, but not enzymatically. If so, we would expect COs in *Exo1^-/-^* spermatocytes to share features with COs from a nuclease-dead allele of *Mlh3* (*Mlh3^DN/DN^*) that replaces a conserved active site residue (D1185N) and eliminates MutLgamma endonuclease activity in vivo and vitro (Nishant et al., 2008; Raghavan, 2019; Toledo et al., 2019).

In *Mlh3^DN/DN^* spermatocytes, MLH1, MLH3, CDK2, and HEI10 form foci with similar timing, frequency, and distribution as WT (Toledo et al., 2019). However, like *Exo1^-/-^*, most of these MutLgamma associated sites fail to become COs. We saw that *Mlh3^DN/DN^* spermatocytes had 5.5 ± 1.9 bivalents per cell on average, like *Exo1^-/-^* (5.3 ± 2.0), but significantly more than *Mlh3^-/-^* (1.4 ± 1.1) (**Figure 3**). Additionally, bivalent distribution was similar between *Mlh3^DN/DN^* and *Exo1^-/-^* with CoVs of 34.7% and 39.1%, respectively. MutLgamma containing *Mlh3^DN/DN^* lacks endonuclease activity (Nishant et al., 2008; Raghavan, 2019; Toledo et al., 2019) suggesting the residual COs seen here are likely derived from SSNs. Consistently, it was previously reported that chiasmata number in *Mlh3^-/-^* and *Mlh3^DN/DN^* spermatocytes partially require MUS81, one of several SSNs (Holloway et al., 2008; Toledo et al., 2019). We find that spermatocytes lacking MUS81 are fertile and show a WT level of bivalents as expected, but that *Mus81^-/-^Exo1^-/-^* spermatocytes had only 4.1 ± 1.9 bivalents per cell, significantly fewer than *Exo1^-/-^*, providing support for the model that the residual COs in *Exo1^-/-^* and *Mlh3^DN/DN^* are derived from SSNs. Taken together, we suggest that residual COs in *Exo1^-/-^* may be independent of MutLgamma endonuclease activity, but dependent upon MutLgamma structurally. In support of MLH3’s structural role, the number of bivalents in *Mlh3^DN^* were dosage sensitive as we saw fewer bivalents per cell on average in *Mlh3^-/DN^* (3.97 ± 1.9) as compared to *Mlh3^DN/DN^*spermatocytes (**Figure 3**). Dosage sensitivity is more consistent with a structural rather than an enzymatic function. Further, we suggest that this structural role of MLH3 must be upstream of EXO1 given the epistatic relationship between *Mlh3^-/-^* and *Exo1^-/-^*.

**Figure 3.**
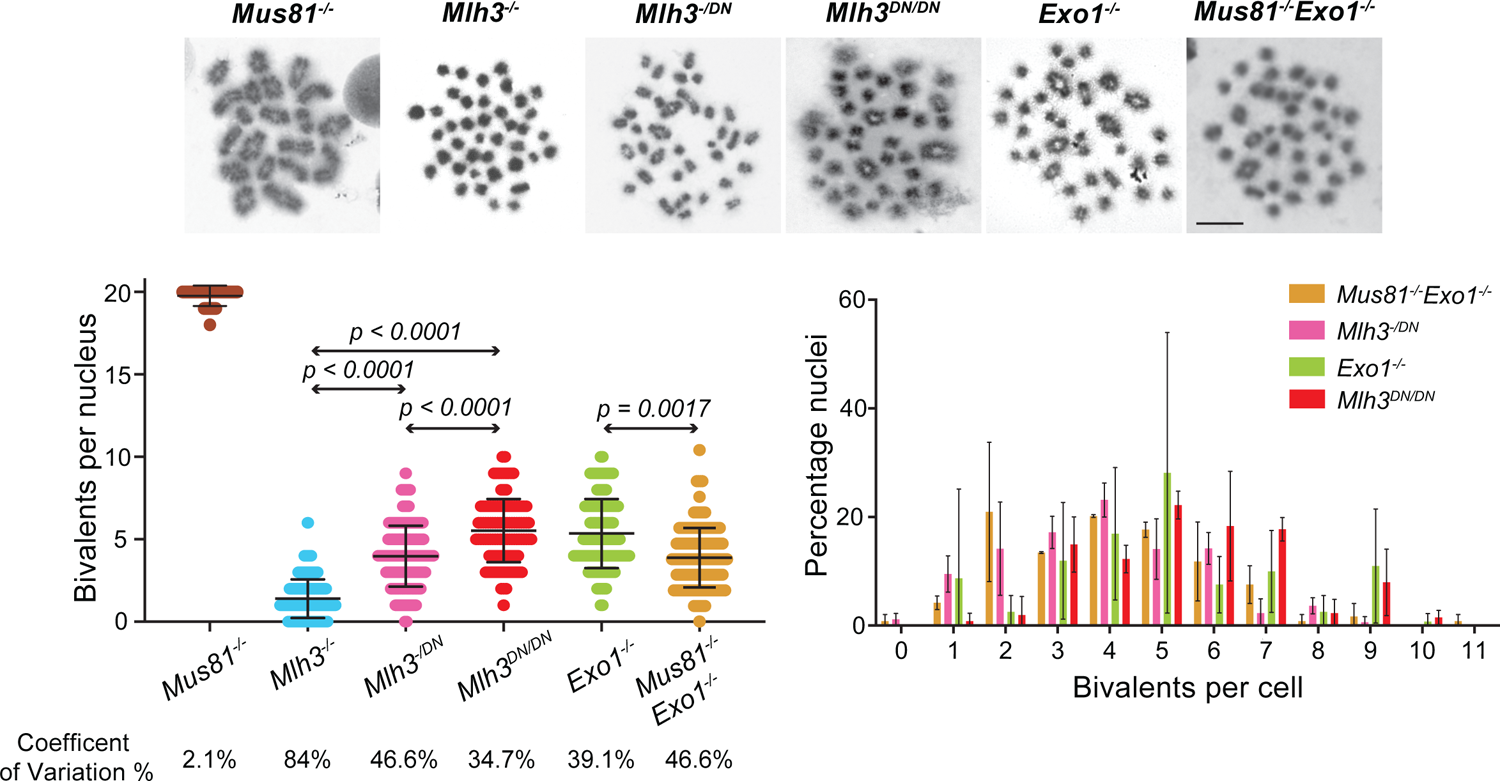
Spermatocytes lacking EXO1 or MLH3 nuclease activity have similar frequency and distribution of bivalents. Representative metaphase images of spermatocytes from the indicated genotypes stained with Giemsa. Left, scatter plot of bivalent counts per cell (y-axis) with error bars of the mean ± SD in *Mus81^-/-^* (N=5, 19.8 ± 0.6), *Mlh3^-/-^* (N=5, 1.4 ± 1.2), *Mlh3^-/DN^* (N=3, 3.97 ± 1.9), *Mlh3^DN/DN^* (N=3, 5.5 ± 1.9), *Exo1^-/-^* (N=4, 5.3 ± 2.0), and *Mus81^-/-^Exo1^-/-^* (N=3, 4.1 ± 1.9). P-values were determined by Kruskal-Wallis test with Dunn’s multiple comparison correction. Right, histogram of the distribution of bivalents per cell of the indicated genotypes. The x-axis shows the number of bivalents per cell and the y-axis shows the average percentage of total nuclei with the indicated number of bivalents. The two graphs also visually depict the CoV of bivalents per genotype: *Mus81^-/-^Exo1^-/-^* (47.8%), *Mlh3^-/DN^* (46.6%), *Exo1^-/-^* (39.1%), and *Mlh3^DN/DN^* (34.7%). *Mlh3^DN/DN^* showed similar frequency and distribution of bivalents as *Exo1^-/-^* spermatocytes, suggesting COs in *Exo1^-/-^* may be independent of MLH3 nuclease activity. Consistently, the number of bivalents was dosage-sensitive to the copy number of *Mlh3^DN^* supporting a structural role for MLH3. Further, loss of the SSN MUS81 in addition to EXO1 (*Mus81^-/-^Exo1^-/-^*) led to fewer bivalents than *Exo1^-/-^* spermatocytes, suggesting some residual COs in *Exo1^-/-^* are formed by SSNs.

To determine when recombination outcomes form in these CO-defective mutants, we synchronized spermatogenesis to enrich for specific stages of meiotic prophase (**Figure 4A**, see **Table S1** for sample purity). We found that in WT, MLH3-derived COs appear during pachynema, while in *Mlh3^-/-^*, SSN-derived COs appear during diplonema (**Figure 4B**). Given the temporal separation between MLH3-dependent and -independent COs, we sought to determine when the residual *Exo1^-/-^* COs appear. We saw that 60% of COs in *Exo1^-/-^* spermatocytes are detectible in diplonema, suggesting the majority of residual COs derive from SSNs (**Figure 4B**). Additionally, we find that residual CO levels at *59.5* are similar between *Mlh3^DN/DN^* and *Exo1^-/-^* Late 4C and diplotene spermatocytes (19.9 ± 9.0, 25.6 ± 7.3, and 22.7 ± 10.8, respectively) (**Table 1**), suggesting the residual COs in *Exo1^-/-^* may be independent of MLH3 nuclease activity. While it is possible that *Mlh3^DN/DN^* and *Exo1^-/-^* have similar numbers of residual COs by coincidence, MutLgamma COs fully require Exo1 in budding yeast (Argueso et al., 2004; Gioia et al., 2021; Nishant et al., 2008; Zakharyevich et al., 2010). We favor a more cohesive model that most residual COs in *Exo1^-/-^*are derived from SSNs and that MutLgamma nuclease activity is heavily reliant on EXO1 in mouse spermatocytes.

**Figure 4.**
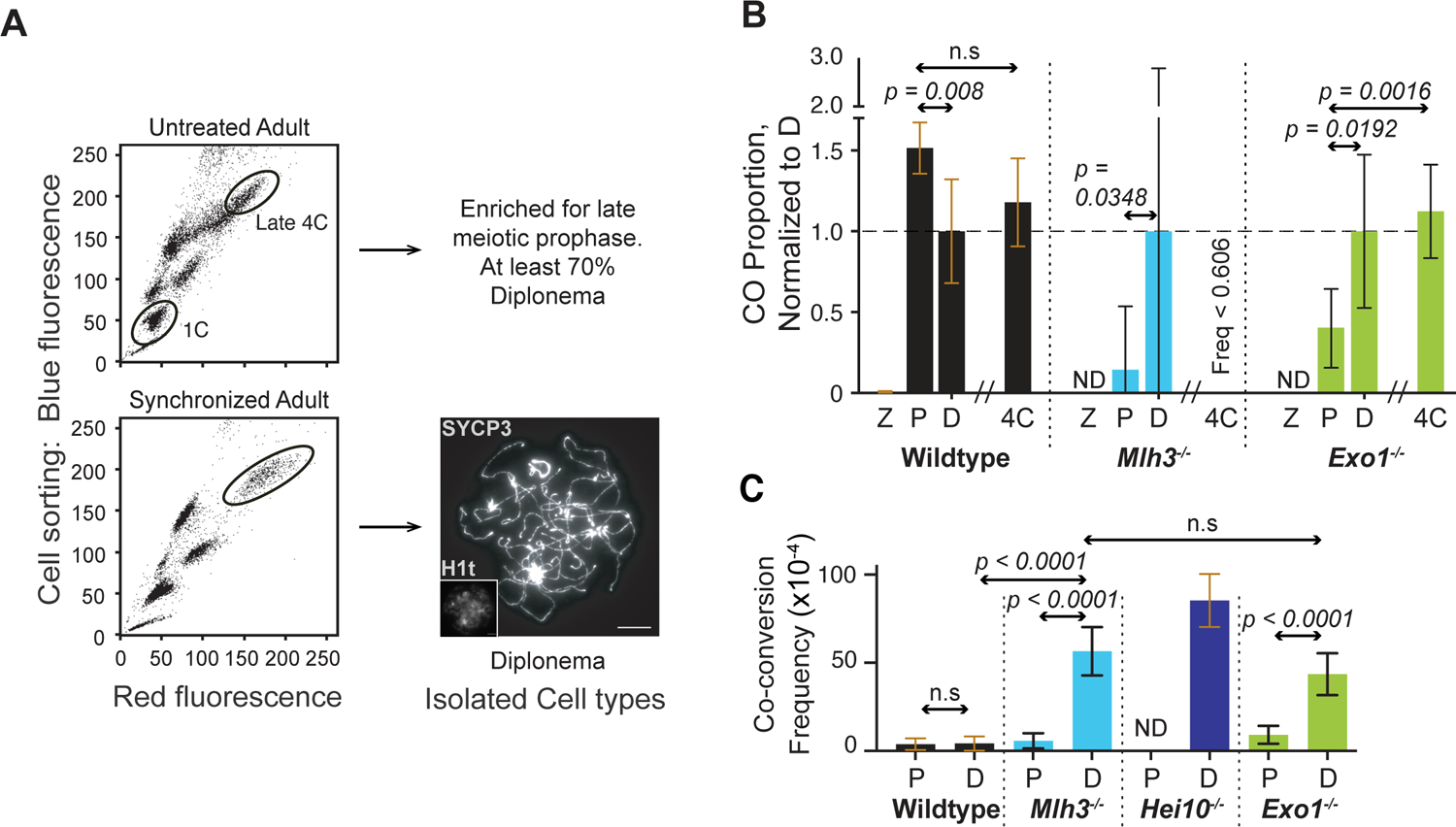
Most COs in *Exo1^-/-^* spermatocytes form contemporaneously with SSN-dependent resolvase activity. **(A)** Top, isolated live testicular cells from an adult BxD mouse were stained with Hoechst 33342 sorted for red and blue emission by UV stimulated fluorescence activated cell sorting. The top oval shows the Late 4C population and sorting gate used to enrich for diplonema and metaphase. The bottom oval shows the population of 1C spermatids. Sorted samples were typically ≥70% enriched for diplonema (**Table S1**). Purity of the sorted population was assessed by immunofluorescence staining of spermatocyte spreads with antibodies against SYCP3 and SYCP1 or gammaH2AX. Bottom, sorted cells as above from an adult BxD mouse that was synchronized for meiotic entry. The example here depicts sorting for a synchronized diplotene population. Quality of synchronization was assessed by pre-sort spermatocyte spreads stained with SYCP3, SYCP1, gammaH2AX, and Histone H1t (**Table S1**). ≥90% of enrichment for specific prophase stages can be achieved by coupling synchronization with sorting. Enrichment of sorted cells was similarly tested by immunofluorescence staining. A representative image of a sorted diplotene stage spermatocyte is shown stained for SYCP3 and Histone H1t (H1t, inset). **(B)** Histogram of the proportion of COs (normalized to the diplonema frequency) found in Late 4C and enriched populations of the indicated stages of meiotic prophase from WT, *Mlh3^-/-^*, and *Exo1^-/-^*. P-values were determined by Fisher’s exact test, two-tailed. In WT, most COs appeared during pachynema with no further increase in diplonema consistent with the model that MutLgamma produces COs in pachynema. By contrast, in *Mlh3^-/-^*, COs in pachynema account for only 14.3 ± 39.2% of COs found in diplonema, consistent with the model that SSN activity is retricted to either diplonema or the pachynema to diplonema transition, likely acting in a back-up repair capacity. The temporal dynamics of *Exo1^-/-^* COs is like that of *Mlh3^-/-^*with COs in pachynema accounting for only 39.9 ± 24.4 % of COs in diplonema. Together, the data suggest that most COs in *Exo1^-/-^* spermatocytes are produced at a stage consistent with SSN activity. **(C)** Histogram of co-conversion frequency found in populations enriched for the indicated stages of meiosis in WT, *Mlh3^-/-^*, *Hei10^-/-^*, and *Exo1^-/-^*. When CO formation is defective, long co-conversions are formed in lieu of COs. Co-conversions are particularly apparent at hotspots like *59.5* that are highly enriched for crossing over. In WT, co-conversions are found at a low frequency in both pachynema and diplonema (3.8 ± 3.3 and 4.2 ± 4, respectively). By contrast, in *Mlh3^-/-^*, co-conversions are similarly low in pachynema but highly enriched in diplonema (5.7 ± 4.3 and 56.5 ± 13.7, respectively). Co-conversions are also highly prevalent in *Hei10^-/-^* diplonema (85.3 ± 15.0), consistent with the model that long co-conversions are derived from RNF212-designated CO precursors. Like *Mlh3^-/-^*, in *Exo1^-/-^* spermatocytes, co-conversions are low in pachynema and highly enriched in diplonema (9.1 ± 5.1 and 40.4 ± 11.4, respectively). Together, most co-conversions found in *Exo1^-/-^* and *Mlh3^-/-^* spermatocytes are likely products of backup repair pathways, acting on unrepaired CO intermediates in either the pachynema to diplonema transition or in diplonema. P-values were determined by Fisher’s exact test, two-tailed. Zygonema (Z), Pachynema (P), Diplonema (D), Late 4C (4C).

### Alternative DNA repair outcomes in *Exo1^-/-^* spermatocytes appear in diplonema

Due to the preponderance of COs at *59.5*, we observe a high frequency of co-conversions in spermatocytes defective for CO maturation, such as *Mlh3^-/-^*and *Hei10^-/-^* (**Figure S1B, Table 1**). We have previously shown that in *Mlh3^-/-^* spermatocytes, these co-conversions likely arise via SDSA and/or dissolution of CO designated intermediates (Zelazowski et al., 2017). In support of this model, we only see enrichment of co-conversions on the B (recipient) chromosome on which we find most CO-dependent gene conversion and never on the D (donor) chromosome as would be expected from SSN-dependent resolution (Marsolier-Kergoat et al., 2018). Finally, we see no enrichment of co-conversions in mutants lacking CO designation such as *Rnf212^-/-^* (**Figure S1B** and L.P, F.C unpublished data). To determine when these co-conversions form, we examined recombination in synchronized *Exo1^-/-^* compared to WT and *Mlh3^-/-^*spermatocytes (**Figure 4C, Table 1**). In WT, co-conversions form at a low frequency in pachynema and we do not see an increase in diplonema. *Mlh3^-/-^* and *Exo1^-/-^* spermatocytes have similarly low frequencies of co-conversions in pachynema as WT frequencies at all stages. Whereas in diplonema, *Mlh3^-/-^* spermatocytes had 10-fold more co-conversions than in pachynema, as well as 13- and 6-fold more co-conversions as WT diplonema and sperm, respectively (**Figure 4C**, **5A**). Similarly, *Exo1^-/-^*had 5-fold more co-conversions in diplonema than pachynema, and 10- and 5-fold more co-conversions in diplonema than WT spermatocytes and sperm, respectively. We suggest that the co-conversions in *Exo1^-/-^* and *Mlh3^-/-^* spermatocytes are products of back-up repair pathways acting on unrepaired CO-designated intermediates in diplonema or during the pachynema to diplonema transition. Intriguingly, despite the high levels of SDSA leading to NCOs in pachynema (**Table 1**), CO intermediates appear inaccessible to SDSA. We find that if CO intermediates are designated, they can only be resolved by MutLgamma in pachynema or repaired by back-up pathways later.

**Figure 5.**
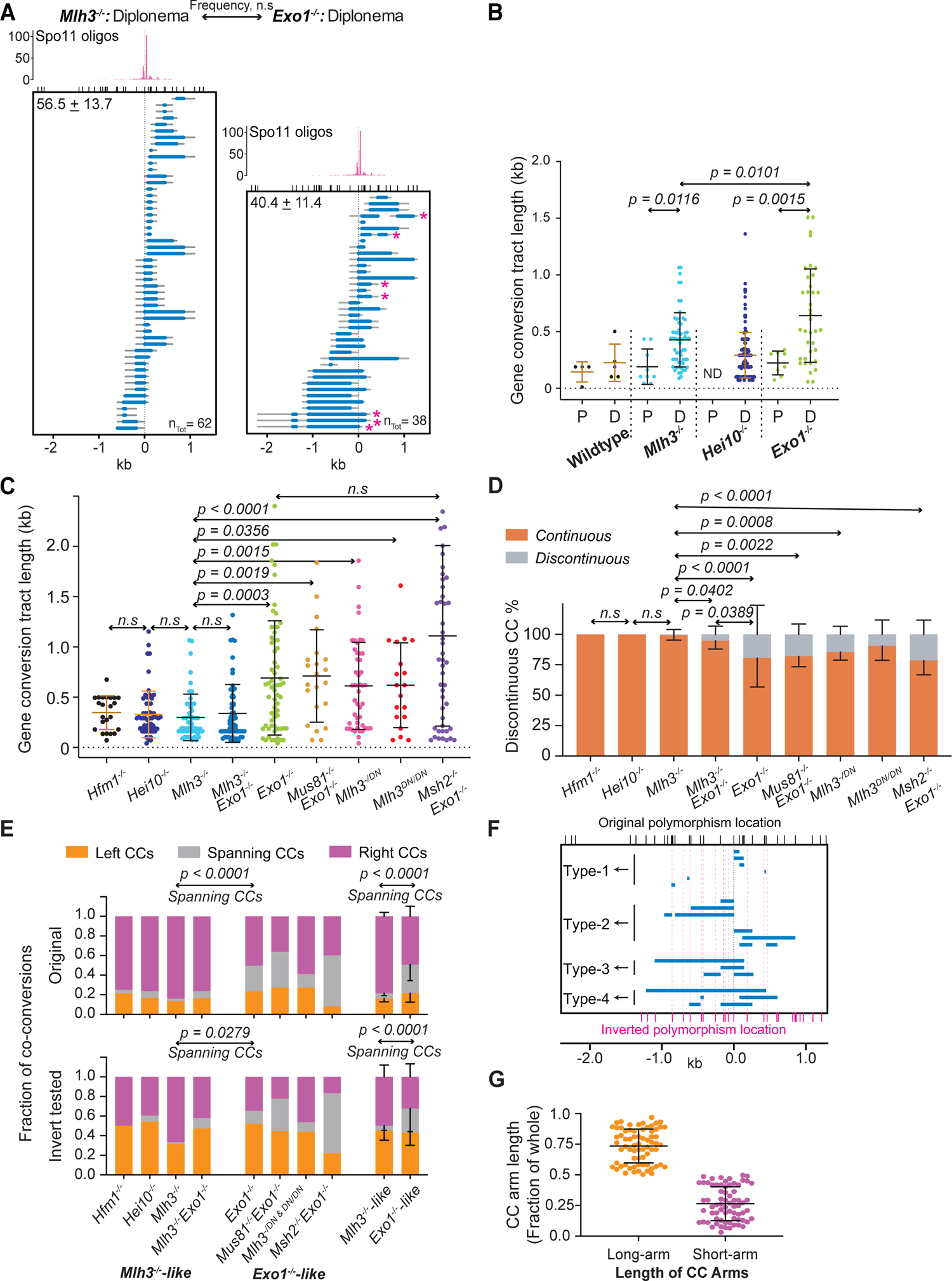
Co-conversion characteristics reveal features of CO precursor processing in mouse spermatocytes. **(A)** Representative population of co-conversions isolated from *Mlh3^-/-^* and *Exo1^-/-^*diplotene spermatocytes. Top, Spo11 oligonucleotides mapping in reads per million in B (Lange et al., 2016) shows the majority of DSBs are formed in the central 200bp of *59.5*. The blue line plots the minimal and the gray line the maximal possible gene conversion tract. The x-axis shows ticks marking polymorphisms at the top and relative position within the hotspot at the bottom. The center of the hotspot is marked by a vertical dotted line and plotted at 0kb. Total frequency ± SD (top left) and number isolated (n_Tot_, bottom right) of co-conversions are shown. The number of co-conversions plotted is proportional to their frequency for visual comparison between genotypes. Asterisks mark discontinuous events. P-value was determined by Fisher’s exact test, two-tailed. While the frequency of co-conversion is similar between *Mlh3^-/-^* and *Exo1^-/-^*, the characteristics are markedly different with *Exo1^-/-^* spermatocytes having longer, discontinuous tracts that frequently span the hotspot center (see below). **(B)** Scatter plot of the length of co-conversions in kb (mean ± SD) in pachynema and diplonema in WT (145 ± 89, 226 ± 164bp, respectively), *Mlh3^-/-^* (190 ± 157, 427 ± 239bp), *Hei10^-/-^* (ND, 292 ± 200bp), and *Exo1^-/-^* (223 ± 105bp, 813 ± 511bp). P-values are derived from Kruskal-Wallis test with Dunn’s multiple comparison correction. In *Mlh3^-/-^*and *Exo1^-/-^* spermatocytes, co-conversions were longer in diplonema than pachynema, consistent with the derivation of long co-conversions from back-up repair pathways acting later. Further, the diplotene stage co-conversions from *Exo1^-/-^* were longer than those from *Mlh3^-^*, suggesting that the upstream CO precursor from which co-conversions are derived is longer in *Exo1^-/-^* than *Mlh3^-/-^* spermatocytes. **(C)** Scatter plot of the length of co-conversions in kb (mean ± SD) in Late 4C from *Hfm1^-/-^*(347 ± 167bp), *Hei10^-/-^* (329 ± 231bp), *Mlh3^-/-^* (298 ± 230bp), *Mlh3^-/-^Exo1^-/-^* (334 ± 288bp), *Exo1^-/-^* (692 ± 567bp), *Mus81^-/-^Exo1^-/-^* (711 ± 459bp), *Mlh3^-/DN^* (613 ± 432bp), *Mlh3^DN/DN^* (619 ± 422bp), and *Msh2^-/-^Exo1^-/-^* (1111 ± 898bp). P-values are derived from Kruskal-Wallis test with Dunn’s multiple comparison correction. The longer co-conversions in *Exo1^-/-^*spermatocytes require MLH3. *Hfm1^-/-^*, *Hei10^-/-^*, and *Mlh3^-/-^* have co-conversions with similar lengths, suggesting mutants that lack MutLgamma focus formation do not extend the CO precursor intermediate. Co-conversions from *Msh2^-/-^Exo1^-/-^* and *Mus81^-/-^Exo1^-/-^* have similar length to those from *Exo1^-/-^*, suggesting neither defective mismatch repair nor SSN-dependent resolution contributes to longer co-conversions. In support of the model that MutLgamma focus formation correlates with longer co-conversions and by extension longer CO precursors, co-conversions from both *Mlh3^-/DN^* and *Mlh3^DN/DN^* were similarly long as those isolated from *Exo1^-/-^* spermatocytes. **(D)** Histogram of the percent of continuous (salmon) and discontinuous (gray) co-conversions (CC) of the indicated genotypes. Discontinuity percentage: *Hfm1^-/-^* (0 ± 0%), *Hei10^-/-^* (0 ± 0%), *Mlh3^-/-^* (0.8 ± 3.9%), *Mlh3^-/-^Exo1^-/-^* (5.3 ± 6.8%), *Exo1^-/-^*(19.4 ± 23.9%), *Mus81^-/-^Exo1^-/-^* (18.1 ± 8.5%), *Mlh3^-/DN^* (14.6 ± 6.6%), *Mlh3^DN/DN^* (9.5 ± 11.8%), and *Msh2^-/-^Exo1^-/-^* (21.5 ± 11.7%). Co-conversions from diplonema and Late 4C spermatocytes were compiled and p-values were determined by Fisher’s exact test, two-tailed. Co-conversions were considered discontinuous when there were two or more contiguous tracts in the same sample, as these were present at a higher frequency than expected by chance. Co-conversions from *Hfm1^-/-^* and *Hei10^-/-^* had similarly low frequency of discontinuity as *Mlh3^-/-^* spermatocytes, whereas *Exo1^-/-^*, *Mus81^-/-^Exo1^-/-^*, *Mlh3^-/DN^*, and *Msh2^-/-^Exo1^-/-^* were more frequently discontinuous. Co-conversions from *Mlh3^-/-^ Exo1^-/-^*spermatocytes were more discontinuous than *Mlh3^-/-^* but less than *Exo1^-/-^*. Discontinuity of co-conversions is the only phenotype where *Mlh3* is not epistatic to *Exo1* suggesting discontinuity is primarily driven by template switching during elongation of the CO precursor and is partially caused by defective MMR. **(E)** Histogram of the distribution of the minimum gene conversion tracts of co-conversions (CC) with respect to the hotspot center. P-values were determined by Fisher’s exact test, two-tailed. Top, all co-conversions. Bottom, to correct for differences in polymorphism density between the left and right side of the hotspot center, polymorphism locations were mirror-inverted with respect to co-conversions, and only those that would have remained detectable were included. No bias in 3’ end invasion of DSBs were observed after correcting for polymorphism density. *Exo1^-/-^-like* mutants had more co-conversions that span the hotspot center than those from *Mlh3^-/-^-like* mutants. We suggest that in *Mlh3^-/-^-like* mutants polymerization occurs from only one of the 3’ ends, whereas in *Exo1^-/-^-like* mutants extension occurs from both 3’ ends. **(F)** Representative co-conversions and their corresponding dispensation after mirror-inverting polymorphism positions. Bottom, relative position within the *59.5* hotspot. Black ticks (top) mark the original polymorphism locations and magenta vertical doted lines and ticks (bottom) mark the mirror-inverted polymorphism locations. The hotspot center is indicated with a black vertical line and mapped at ‘0”. Co-conversions were categorized into 4 groups: Type-1) one-sided co-conversions that would not have been detected, Type-2) one-sided co-conversions that would remain detected, Type-3) co-conversions that span the hotspot center originally, but would be detected as one-sided with inverted polymorphisms, and Type-4) co-conversions that span the hotspot center originally and would remain detected as spanning after inverting polymorphisms. **(G)** Scatter plot showing the fraction of each gene conversion tract contributing to either the long or short arm of the co-conversions (CC) that span the hotspot center. This marked skew in the distribution of co-conversions is likely caused by differential polymerization from the independent 3’ ends.

### Co-conversion length in CO defective mutants suggests additional steps in CO intermediate processing

Having established that *Exo1^-/-^* co-conversions at *59.5* likely arise from back-up repair pathways acting after pachynema, we sought to define the features of these co-conversions. The lengths, degree of discontinuity, genetic dependence, and distribution of these co-conversions should reflect characteristics of the CO designated intermediates from which they are derived providing mechanistic insight. For example, co-conversions that form during diplonema in CO maturation mutants should be longer than those that arise via SDSA in pachynema. Consistent with this concept, we saw that co-conversions in *Mlh3^-/-^* and *Exo1^-/-^* pachytene spermatocytes were 190 ± 157bp and 223 ± 105bp long, respectively (**Figure 5B**, R.K., F.C. unpublished data) and were similar in length to each other and those isolated from WT pachynema and diplonema, suggesting that they arise from similar intermediates and likely via SDSA. By contrast, co-conversions detected in *Mlh3^-/-^*and *Exo1^-/-^*diplotene spermatocytes averaged 427 ± 239bp and 813 ± 511bp, respectively, and were significantly longer than co-conversions from their pachytene counterparts or WT spermatocytes. Curiously, we saw that co-conversions from *Exo1^-/-^* diplotene spermatocytes were significantly longer than those from *Mlh3^-/-^* (**Figure 5A, B**). Considering that the co-conversion tract length likely reflects its precursor intermediate, we suggest that the CO designated precursor elongates in *Exo1^-/-^*, a process that may require MLH3.

### Elongated co-conversions in *Exo1^-/-^* spermatocytes require MLH3 structurally

To definitively determine the genetic dependence of the frequency, distribution, and length of co-conversions, we analyzed NCOs in multiple CO maturation mutants and double mutants. Similar to observations in diplonema, we saw that co-conversions isolated from Late 4C *Exo1^-/-^* spermatocytes were longer (692 ± 567bp) than those isolated from *Mlh3^-/-^* (298 ± 231bp) (**Figure 5C**). We also saw that co-converted NCOs from *Exo1^-/-^Mlh3^-/-^* spermatocytes were similar in length (334 ± 288bp) to those isolated from *Mlh3^-/-^*. As such, MLH3 is required for the elongated co-conversions seen in the absence of EXO1. Generally, *Exo1^-/-^* has a stronger mismatch correction defect than MutLgamma (Furman et al., 2021), likely ruling out defective mismatch repair as a significant contributor to elongation of the co-conversions seen in *Exo1^-/-^* spermatocytes. Further supporting this premise, we found that co-conversions isolated from *Exo1^-/-^Msh2^-/-^* were not significantly changed in length (1111 ± 898bp) to those from *Exo1^-/-^* spermatocytes. Elongation of co-conversions in *Exo1^-/-^* spermatocytes was independent of MUS81, supporting our model that co-conversions likely result from dissolution or SDSA-like mechanisms, rather than from double Holliday junction resolution, although we cannot rule out that redundant SSN pathways play a role. We suggest elongation of the designated CO precursor leads to elongation of the co-conversions in *Exo1^-/-^*, *Exo1^-/-^Msh2^-/-^*, and *Exo1^-/-^Mus81^-/-^* as compared to *Mlh3^-/-^* or *Exo1^-/-^Mlh3^-/-^* spermatocytes.

A major difference between the mutants with shorter and longer co-conversions is that MutLgamma accumulates at CO sites in *Exo1^-/-^*, *Exo1^-/-^Msh2^-/-^*, and *Exo1^-/-^Mus81^-/-^* spermatocytes (**Figure 1C)** (Kan et al., 2008; Wei et al., 2003). Therefore, we sought to test whether MutLgamma focus formation is required to observe elongation of co-conversions and of the inferred designated CO intermediate. We analyzed NCO frequency and distribution in *Hei10^-/-^* and *Hfm1^-/-^* spermatocytes. In mouse spermatocytes, HEI10 is proposed to target MutSgamma for ubiquitination and degradation allowing exchange in favor of MutLgamma at designated CO sites (Qiao et al., 2014; Rao et al., 2017; Ward et al., 2007). HFM1 is a Mer3 helicase ortholog and is required for full synapsis, formation of MutLgamma foci at CO sites, and 80% of chiasmata in mouse spermatocytes (Guiraldelli et al., 2013). As such, MLH1 and MLH3 are present in *Hei10^-/-^* and *Hfm1^-/-^* spermatocytes, but do not form MutLgamma foci on CO sites. In *Hfm1^-/-^* and *Hei10^-/-^*, co-conversions were similar in length (347 ± 167bp, 329 ± 231bp, respectively) to co-conversions from Late 4C *Mlh3^-/-^* spermatocytes (**Figure 5C**). We infer that the presence of MLH3 alone is insufficient to elongate the co-conversions. Additionally, the similar level of co-conversions found in *Hfm1^-/-^* compared to *Mlh3^-/-^* spermatocytes suggests that unlike yeast Mer3 (Borner et al., 2004), HFM1 is not required for CO designation.

To test if MutLgamma focus formation correlates with longer co-conversions, we analyzed *Mlh3^DN/DN^* spermatocytes. Similar to *Exo1^-/-^*, the *Mlh3^DN/DN^* allele is defective for CO maturation, but is proficient for MutLgamma focus formation at likely designated CO sites (Toledo et al., 2019). Interestingly, we observed that co-conversions from *Mlh3^DN/DN^* and *Mlh3^-/DN^* (619 ± 422bp and 613 ± 432bp, respectively) were similar in length to co-conversions from *Exo1^-/-^* and significantly longer than those from *Mlh3^-/-^* spermatocytes (**Figure 5C**). Indirect estimations and direct measurements show that CO gene conversion tract lengths in mice and humans are ∼600 bp at multiple hotspots (Cole et al., 2014; Jeffreys and Neumann, 2002; Zelazowski et al., 2017). Comparing the co-conversions across multiple mutants, we observe that those with MutLgamma foci have longer gene conversion tracts than those without, and that the longer tracts resemble mammalian CO-dependent gene conversion (**Figure 5C**). We suggest that the CO precursor intermediate has further differentiated in mutants that load MutLgamma on the axes. In support of this concept, there are more SSN-derived bivalents in *Exo1^-/-^* and *Mlh3^DN/DN^* suggesting the precursor intermediate may have progressed further towards CO maturation capability (**Figure 3**). We suggest this progression also occurs in wild type upon association of MutLgamma, but is rapidly resolved as a CO because MLH3 is proficient for nuclease activity. Taken together, the inferred elongation of the designated CO intermediate is associated with MutLgamma focus formation, independent of MLH3 nuclease activity, and correlates with increased probability of resolution as a CO.

### Co-conversions from *Exo1^-/-^* and *Mlh3^DN/DN^* are frequently discontinuous

Discontinuous co-conversions involve a single event that has four genotype switches between parental haplotypes (e.g., asterisks, **Figure 5A**). An apparent discontinuous co-conversion could arise from two independent NCOs occurring in the same samples. Indeed, the number of discontinuous co-conversions that involved a singleton and a contiguous co-conversion were found at the frequency predicted by Poisson approximation for multiple events per sample. As such, these events were removed from the analysis (Supplementary data file). By contrast, the number of discontinuous co-conversions that involved two separate contiguous co-converted tracts were much higher than predicted by Poisson. On average a greater fraction of co-conversions were discontinuous in *Exo1^-/-^* (19.4 ± 23.9%) as compared to *Mlh3^-/-^* spermatocytes (0.8 ± 3.9%) (**Figure 5D**). This 24-fold higher frequency of discontinuity cannot be explained by the ∼2-fold longer co-conversion tract length between *Mlh3^-/-^* and *Exo1^-/-^* spermatocytes. Discontinuity may involve defects in mismatch repair, as expected based on the role of EXO1 in MLH1-dependent MutLalpha activity. Consistent with this premise, levels of discontinuity were similar between *Exo1^-/-^* and *Msh2^-/-^Exo1^-/-^* (21.5 ± 11.7%), suggesting absence of EXO1 alone is sufficient for this phenotype. Loss of MUS81 does not influence the frequency or discontinuity of co-conversions consistent with their formation likely arising from dissolution and SDSA-like mechanisms. Interestingly, we saw that *Mlh3^-/-^Exo1^-/-^* has an intermediate phenotype with the fraction of discontinuous co-conversions (5.3 ± 6.8%) being significantly different from both *Mlh3^-/-^* and *Exo1^-/-^* (**Figure 5D**). Discontinuity of co-conversions is the only phenotype for which *Mlh3* is not epistatic to *Exo1*, supporting the interpretation that mismatch repair plays at least a partial role in formation of discontinuos co-conversions. Not exclusively, discontinuity could reflect template switching in the formation of the elongated CO intermediate. In support of this model, co-conversions in *Mlh3^-/DN^* had a high fraction of discontinuity (14.6 ± 6.6%), suggesting either template switching and/or lack of efficient mismatch repair. Similarly, co-conversions in *Mlh3^DN/DN^* were also discontinuous (9.5 ± 11.8 %), however there were too few events for analysis. Perhaps the presence of nuclease-dead MLH3 may impede MLH1 function in MutLalpha mismatch repair in this context. However, template switching is frequently observed in budding yeast lacking MLH3 or EXO1 (Marsolier-Kergoat et al., 2018), and has been proposed to occur frequently during CO formation (Ahuja et al., 2021). We favor the model that defects in mismatch repair contribute to discontinuity, but the lion’s share is owing to template switching upon elongation of the designated CO intermediate. In support of this model, we find a similar fraction of discontinuos COs and co-conversions in *Exo1^-/-^* (**Figure 2B**). Further, we see a two-fold higher frequency of discontinuous COs in *Exo1^-/-^* than in *Exo1^nd/nd^*spermatocytes, suggesting additional mechanisms contribute to discontinuity than defective mismatch repair. Importantly, when MLH3 retains its EXO1-dependent nuclease activity, discontinuity is not apparent, suggesting that the MutLgamma nicking to resolve COs also contributes to conversion or reversion of mismatches as predicted in yeast and previously suggested in mouse spermatocytes (Peterson et al., 2020).

### Both 3’ ends of a DSB are equally likely to invade and polymerize from the homolog at *59.5*

In order to glean any evidence for how the CO-designated intermediate is formed and differentiated, we analyzed the distribution of co-conversions in the two classes of CO maturation mutants (*Mlh3^-/-^*-like and *Exo1^-/-^*-like). Importantly, *59.5* is an asymmetric hotspot in the BxD F1 hybrid and experiences a higher frequency of SPO11-dependent DSBs on the B rather than the D chromosome. The majority of these DSBs occur directly at the hotspot center (**Figure 5A**) (Lange et al., 2016) (marked with a dashed line and plotted at 0kb throughout), where a polymorphism confers a high affinity PRDM9 binding site on the B chromosome that specifies the *59.5* hotspot. Consistent with the differential break frequency between homologs, there is a concomitant three-fold higher frequency of NCOs in WT on the B versus D chromosome and reciprocal CO asymmetry (Zelazowski et al., 2017). Intriguingly, the long co-conversions seen in *Mlh3^-/-^*-like and *Exo1^-/-^*-like mutants are found exclusively on the B chromosome, suggesting that only DSBs initiated on B are capable of becoming designated CO sites. The asymmetry of meiotic DSBs at *59.5* and the subsequent asymmetry of the resulting co-conversions provides an opportunity to investigate whether any strand invasion or extension biases can be inferred about the CO-designated intermediate based upon the distribution of co-conversions.

We categorized the co-conversions as to whether they were to the left, spanning, or to the right of the hospot center (**Figure 5E, top**). In examining the co-conversion distribution, it would appear that there is a marked enrichment for co-conversions on the “right” side of the hotspot center (**Figure 5A**, dotted line). For example, in *Mlh3^-/-^* we find 100 co-conversions to the right and only 16 to the left (**Figure 5E**). This right-sided enrichment was present in all mutants, but was particularly prevalent in mutants of the *Mlh3^-/-^*-like class. This could reflect a preference for strand invasion by one end of the DSB. However, the polymorphism density to the right of the hotspot center is higher than to the left raising the possibility that we have more detectible events on the right side (**Figure 5F, ‘Original polymorphism location’**). To correct for the influence of polymorphism density, we mirror inverted the polymorphism location at the hotspot center from left to right and determined whether the original co-conversions would have been detected given the altered polymorphism map (**Figure 5F, ‘Inverted polymorphism location’**). There were four outcomes after inverting the polymorphisms and examples of each are shown in **Figure 5F**: Type-1) one-sided co-conversions that would not have been detected; Type-2) one-sided co-conversions that would have been detected; Type-3) co-conversions that span the hotspot center originally, but would be detected as one-sided; and Type-4) co-conversions that span the hotspot center originally and would remain detected as such. To counter any potential detection skew, Type-1 co-conversions were removed from the analysis and Type-3 co-conversions were re-scored as one-sided (**Figure 5E, bottom**). Upon this reassessment, we found no significant difference between the frequency of events on either side of the hotspot center, confirming our suspicion that the polymorphism density was driving the apparent right-sided enrichment. In conclusion, either end of the DSB is equally likely to engage in strand invasion at *59.5*.

### Co-conversion distribution suggests step wise polymerization of 3’ invaded ends

One intriguing observation in the *Mlh3^-/-^*-like class of mutants is that only 4% of co-conversions span the hotspot center (**Figure 5A,E**). If both ends of the DSB extend via polymerization, we would expect most of these co-conversions would incorporate the hotspot center, despite their relatively short average length of 311bp. Therefore, we suggest that the CO-designated intermediate in the *Mlh3^-/-^*-like class of mutants is extended from only a single invaded 3’ end. By contrast, 25% of events in the *Exo1^-/-^*-like class of mutants span the hotspot center suggesting that both ends of the DSB are extended. Why then don’t all *Exo1^-/-^*-like co-conversions span the hotspot center? These spanning co-conversions show a marked skew in distribution with ∼72% occuring on only one side of the hotspot resulting in a long and short arm of the co-conversions (**Figure 5G**). We suggest that upon MutLgamma loading in the *Exo1^-/-^*-like class of mutants, the initial invaded 3’ end is further extended by polymerization and the second end is 3-fold less polymerized on average. The shorter extended tract on the second end would frequently fail to incorporate a polymorphism such that the majority of events still appear one-sided. Consistent with this premise, 40% of the spanning co-conversions in the *Exo1^-/-^*-like class were categorized as Type-3 outcomes upon mirror inverting polymorphisms and would have been detected as one-sided (**Figure 5F**).

A consequence of the shorter polymerization of one 3’ end in *Exo1^-/-^* would be that upon resolution of this intermediate, one of the CO breakpoints should be close to the hotspot center, which is what we observe in the CO breakpoint mapping (**Figure 2C**). If we plot the ends of the *Exo1^-/-^* co-conversions as hypothetical CO breakpoints, we find our hypothetical CO breakpoint map is similar to the observed CO break point map in *Exo1^-/-^* with similar reciprocal CO asymmetry and shorter inferred gene conversion tract length (**Figure S2**). Considering the shorter single-end extended CO precursor in the *Mlh3^-/-^*-like mutants, we would expect the SSN dependent COs to form even closer to the hotspot center than in *Exo1^-/-^* spermatocytes. Indeed, we find that 85% of COs in *Mlh3^-/-^*-like mutants (12 out of 14) are found in the central 500bp as opposed to 54% in *Exo1^-/-^* (53 out of 98) and 32% in *Exo1^het^* (30 out of 92) (0.0398 and 0.0002, respectively Fisher’s exact test, two-tailed). Taken together, we suggest the single-end extended intermediate is independent of MLH3, the dual-end extended intermediate is dependent upon MLH3 structurally, and that both are likely formed in pachynema. In the absence of MutLgamma resolvase activity, both single- and dual-end intermediates are substrates that can be resolved by SSNs or dissolved after pachynema.

Unlike *Exo1^-/-^*, in wild type, *Exo1^het^*, and *Exo1^nd^* the majority of COs are simple and their breakpoints show more reciprocal CO asymmetry (**Figure 2B, 2C, S2**). We surmise that when MutLgamma is enzymatically proficient, retained heteroduplex and evidence of template switching is less frequent and CO breakpoints are more evenly distributed away from the hotspot center. Perhaps MLH3-dependent nicking can direct mismatch repair and/or strand processing during CO formation. In support of this model, genomic analysis of COs in *Msh2^-/-^* budding yeast show fewer heteroduplex strands as compared to those lacking Msh2 along with Mlh1, Mlh3, Exo1, or Sgs1(Marsolier-Kergoat et al., 2018). Further, in depth analysis of COs at one highly polymorphic hotspot in *Msh2^-/-^* budding yeast suggests frequent CO resolution-associated strand processing (Ahuja et al., 2021), and retained heteroduplex is rare in *Msh2^-/-^* mouse spermatocytes (Peterson et al., 2020).

Taken together, we suggest that the co-conversion pattern in the *Mlh3^-/-^*-like and *Exo1^-/-^*-like class of mutants reflects two independent DNA repair intermediates (**Figure 6**). First a single end invades and extends ∼300bp to form an extended single-end invasion intermediate (eSEI), this relatively stable intermediate requires MutLgamma to be loaded on the axis for the second end to be captured forming a nicked double Holliday junction. Both ends are extended by polymerization, but the second end polymerizes to a shorter extent. Alternatively, second end capture could occur prior to MutLgamma loading with minimal or no extension occuring on the second end. Finally, MutLgamma nicking directs strand processing and mismatch repair to reorient the gene conversion tract on either side of the hotspot center.

**Figure 6.**
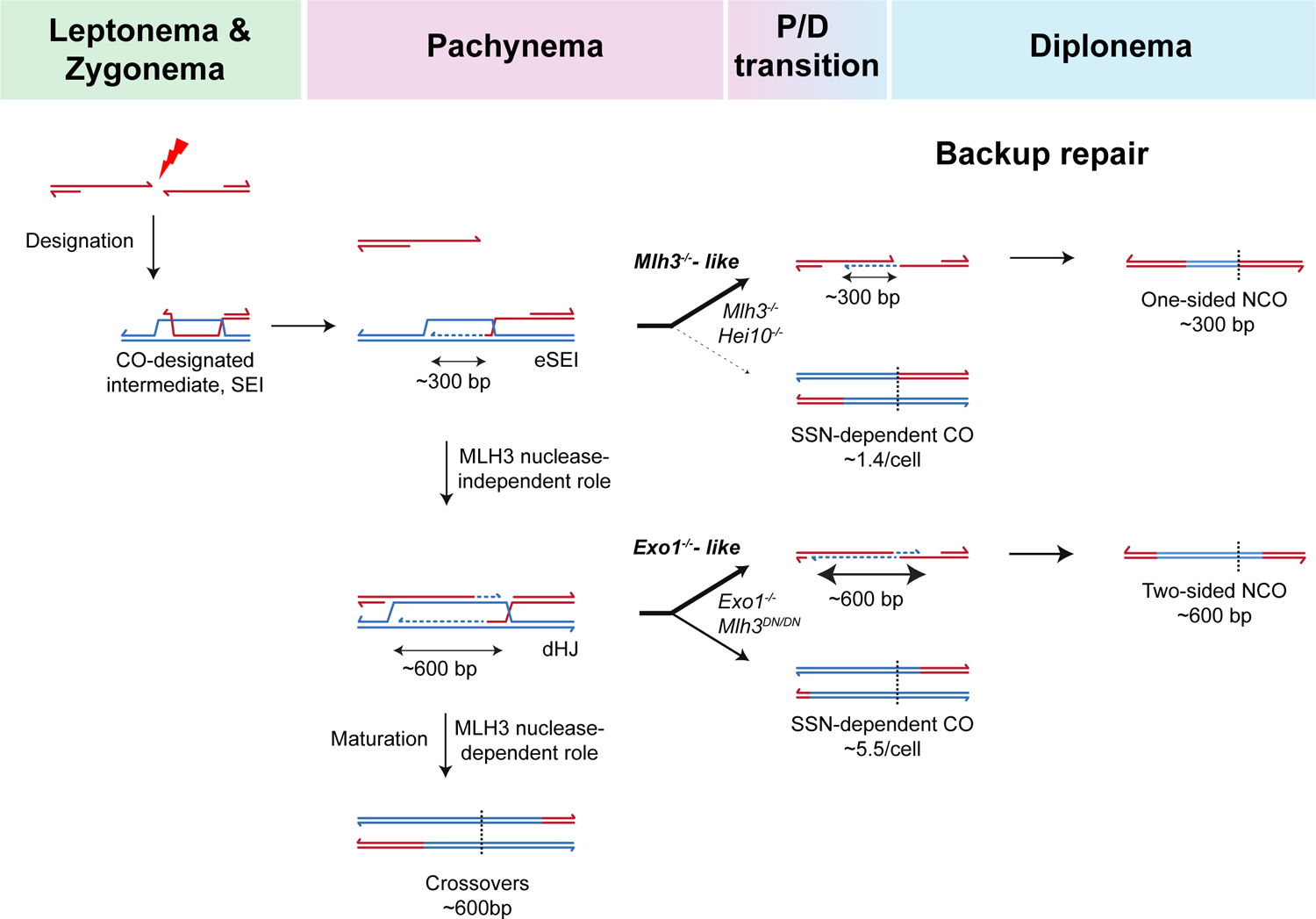
Model illustrating CO precursor differentiation and maturation in mouse spermatocytes. Either 3’ end of the DSB is equally likely to invade the homolog to form the CO-designated intermediate - the single-end invasion (SEI). In pachynema, the invaded 3’ end extends via polymerization for ∼300bp forming an extended SEI (eSEI) intermediate. Formation of the eSEI is MLH3 independent, but invasion/extension of the second 3’ end to generate a nicked double Holliday junction (dHJ) depends upon MLH3’s structural function. This second round of polymerization extends both invaded 3’ ends but asymmetrically, to a total length of ∼600bp. This nicked dHJ is more capable of being resolved as a CO via SSN-mediated resolution. Finally, we suggest that during resolution, nicking by MLH3 directs strand processing and mismatch repair that reorients CO breakpoints symmetrically on either side of the hotspot center with few or no haplotype switches. By contrast, SSN-dependent resolution lacks the ability to reorient CO breakpoints as observed in *Exo1^-/-^*- and *Mlh3^-/-^*-like spermatocytes. The vertical black dotted line represents the location of the originating DSB.

## Conclusions

While many of the meiotic recombination mechanisms are conserved between budding yeast and mammals, there are unique differences that provide opportunities to refine recombination models. To that end, recombination analysis has been done in mouse spermatocytes in WT and a few mutants (*Msh2*, *Mlh1*, and *Mlh3*) (Baudat and de Massy, 2007; Cole et al., 2014; Cole et al., 2010; Guillon et al., 2005; Guillon and de Massy, 2002; Peterson et al., 2020; Svetlanov et al., 2008; Zelazowski et al., 2017). Unexpectedly, mismatch defective mouse spermatocytes retain far less heteroduplex strands as compared to mismatch defective yeast (Ahuja et al., 2021; Marsolier-Kergoat et al., 2018; Peterson et al., 2020). Additionally, physical detection of CO precursors hasn’t been accomplished in mammals. As a result, we are relatively ignorant of the structure of the CO intermediates in mammals. To overcome these limitations, we analyzed ∼1 million haploid genome equivalents from 12 independent genetic conditions at a highly polymorphic native recombination hotspot. We also determined when recombination is executed in wild type and 2 mutants, and together analyzed ∼1600 COs and ∼4400 NCOs. We verify conservation of several aspects of meiotic recombination between yeast and mouse spermatocytes, including several recently discovered features:

1) Similar to previous observations (Kirkpatrick et al., 2000), we provide evidence that EXO1 nuclease activity contributes to mismatch correction during meiotic recombination (**Figure 2B**).
2) EXO1 structurally promotes MLH3-dependent nuclease function and thereby CO formation in mammals (**Figure 2A**) (Yamada et al., 2020; Zhao et al., 2018), as was first established in budding yeast (Argueso et al., 2004; Gioia et al., 2021; Nishant et al., 2008; Zakharyevich et al., 2010) and verified in mouse (Wang et al., 2022).
3) The inferred CO precursor in mutants that lack MutLgamma foci (*Mlh3^-/-^*-like) is asymmetric regarding the two 3’ ends of the DSB, independent of MutLgamma (**Figure 5**), but dependent on the ZMM protein RNF212 (**Figure S1** and R.K., F.C. unpublished data). We suggest this intermediate is likely a single-end invasion intermediate – the first CO-specific precursor that is physically detectible in yeast (Hunter and Kleckner, 2001).
4) COs and co-conversions derived from CO precursors in mutants that lack MLH3’s enzymatic activity (*Exo1^-/-^*-like) frequently display signatures of template switching and/or branch migration (**Figure 2B, 5D**), similar to recent findings based upon heteroduplex retention in *Msh2^-/-^* budding yeast (Ahuja et al., 2021; Marsolier-Kergoat et al., 2018). Further, we saw evidence that MLH3’s nuclease activity directs substantial mismatch correction during CO formation as was previously proposed (Marsolier-Kergoat et al., 2018).
5) We did not see evidence to support the presence of ligated dHJs in pachynema, which should be amplifiable in our assays (**Figure 4C**). While absence of evidence is not evidence of absence, we favor the model that dHJs are likely nicked, which has been proposed by others. Resolution of nicked dHJs is compatible with the biochemical properties of MutLgamma in vitro as well as the heteroduplex patterns and electron microscopy analysis in budding yeast (Bell and Byers, 1983; Cannavo et al., 2020; Crown et al., 2014; Gioia et al., 2021; Kulkarni et al., 2020; Manhart et al., 2017; Marsolier-Kergoat et al., 2018; Peterson et al., 2020; Stahl et al., 2004)

We also elaborate on and provide support for models based upon observations in mouse spermatocytes:

1) In marked contrast to budding yeast, we confirm by cytological and recombination analysis that the CO phenotype in *Exo1^-/-^* spermatocytes is unlike *Mlh3^-/-^* in mouse (Kan et al., 2008; Wei et al., 2003) (**Figure 1, 2A**).
2) There are mild alterations in the frequency of recombination intermediate foci in spermatocytes lacking EXO1 (Kan et al., 2008) (**Figure 1B, C**).
3) We provide evidence that MUS81 contributes to residual CO formation when MutLgamma lacks endonuclease activity, such as in the absence of EXO1 (**Figure 3**), as is known in budding yeast and mouse (Argueso et al., 2004; Gioia et al., 2021; Holloway et al., 2014; Nishant et al., 2008; Toledo et al., 2019).
4) A structural role that promotes SSN activity was proposed for MLH3 based upon the frequency of chiasmata in *Mlh3^DN/DN^*spermatocytes (Toledo et al., 2019). We provide strong support for the model that MLH3 has a nuclease-independent role (**Figure 3, 4**) that results in increased COs derived from SSNs.

Finally, this work has expanded our understanding of mammalian meiotic recombination by inferring two independent CO precursors, their genetic dependence, and identifying unique roles for CO maturation proteins to establish that:

1) In contrast to budding yeast, in mouse spermatocytes, loss of EXO1 does not alter CO interference (**Figure S1C**) nor is the Mer3 ortholog HFM1 required for CO designation (**Figure 5C, D, E and S3**).
2) While like budding yeast, MutLgamma-derived COs require EXO1 structurally, we establish that in mouse spermatocytes, residual COs in *Exo1^-/-^* require MLH3 structurally (**Figure 3**, **4B**). This unexpected structural role of MLH3 is upstream and independent of EXO1.
3) We establish independent genetic requirements for the two 3’ ends of the DSB to polymerize, suggesting stepwise differentiation of CO precursors into dHJs in mouse spermatocytes. Polymerization of the initial invading 3’ end is independent of MutLgamma, while the structural role of MLH3 promotes either second-end capture and/or polymerization from the second 3’ end (**Figure 5**). This in marked contrast to budding yeast, where dHJs form at similar frequencies in the presence or absence of MutLgamma (Zakharyevich et al., 2010; Zakharyevich et al., 2012).
4) MLH3’s structural role likely promotes formation of dHJs with asymmetric polymerization of the two invaded 3’ ends (**Figure 5G**) and MLH3’s nuclease role subsequently generates simple CO break points that are evenly distributed around the hotspot center likely due to nick-directed MMR (**Figure 5D**). Unlike MutLgamma, SSNs do not reposition CO breakpoints upon resolution (**Figure S2**).
5) The similarities within and differences between the *Mlh3^-/-^-like* and *Exo1^-/-^-like* class of mutants suggests that the MutLgamma-independent and MLH3 structure-dependent CO precursors are stable and stereotypic, suggesting the existence of both single-end invasion and nicked-dHJ intermediates in CO formation in mammals (**Figure 5, 6**).

Whole-genome analysis of CO resolution patterns in budding yeast shows that when MutLgamma is proficient, there are two types of resolution biases: CO-biased resolution of dHJs and a likely independent orientation bias for dHJ resolution that generates a single tract of heteroduplex with no intervening fully converted tracts (Pattern 1) (Marsolier-Kergoat et al., 2018). Heteroduplex patterns at specific hotspots suggests that Pattern 1 COs predominate in mouse spermatocytes as well (Peterson et al., 2020). It is likely that these two resolution biases are rooted in independent asymmetries of the CO precursor intermediate. Indeed we find two such asymmetries: 1) a MutLgamma-independent asymmetry, where only one of the two 3’ strands polymerize upon invasion and 2) an MLH3 structure-dependent asymmetry with unequal polymerization of the two invaded strands. Several lines of evidence suggest that CO designated intermediates are biased towards a CO configuration independent of MutLgamma. For example, Yen1, an SSN with no inherent resolution bias in vitro, generates COs with similar frequencies as MutLgamma when activated after CO designation in budding yeast (Arter et al., 2018). It should also be noted that some organisms such as *Caenorhabditis elegans* lack MutLgamma entirely and rely on SSNs for generation of interference-dependent COs (Saito et al., 2013). However, we do not find an increase in SSN-dependent crossing over until after dHJ formation suggesting that the MLH3 structure-dependent asymmetry is the key driver of CO biased resolution in mouse spermatocytes.

In conclusion, the combination of fine-scale recombination and global cytological analysis in CO maturation deficient mouse spermatocytes allowed us to discover both conserved and unique features of meiotic recombination mechanisms to propel our understanding of CO intermediate processing in general.

## Limitations of the study

Recombination analysis in this study is limited to a single locus, the *59.5* hotspot, which is useful for study due to its high frequency of meiotic DSBs on one parental homolog, density of polymorphisms, and strong preference for crossing over. However, it should be noted that behavior of *59.5* may not reflect genome-wide patterns of recombination. We attempted to compare all available locus-specific analysis to global frequencies using cytology, but some features, such as NCOs are indistinguishable cytologically. Additionally, our model of CO precursor differentiation is inferred based upon the end products of recombination in mutants. As such, confirmation will depend upon further studies in wild type and the development of new technologies to detect CO precursors in mammals.

## Figure Legends

**Figure S1. Axis lengths, recombination frequencies, and interference measurements**

**(A)** Scatter plot of the axis lengths per cell measured in microns (mean ± SD) in WT (185.8 ± 23.6) and *Exo1^-/-^* (186.0 ± 24.8). *Exo1^-/-^* spermatocytes had similar axis lengths as WT.

**(B)** Histogram plot of total recombination frequency (per 10,000 haploid genome equivalents) for the indicated genotype categorized into COs (pink), co-conversions (green) and singletons (blue). Total recombination frequencies were similar among WT sperm (244.6), WT diplonema (213.4), and mutants including *Rnf212^-/-^* Late 4C (196), *Hei10^-/-^* diplonema (211.6), *Mlh3^-/-^* diplonema (194.6), and *Exo1^-/-^* diplonema (193.7). P-values were determined by Fisher’s exact test, two-tailed.

**(C)** Plot of mean coefficient of coincidence (CoC) versus inter-interval distance calculated by measuring MLH1 focus distribution of the indicated genotypes. We saw that MLH1 foci from *Exo1^-/-^* (top right, and bottom – scale adjusted) spermatocytes show similar interference as WT. SP (sperm), D (diplonema), 4C (Late 4C).

**Figure S2. Inferred CO breakpoints from co-conversions in *Exo1^-/-^* spermatocytes have similar distribution as *Exo1^-/-^* COs**

Top, cumulative distribution curves of CO breakpoints in the B to D (red) and D to B (blue) orientations from *Exo1^het^*, *Exo1^-/-^*, and inferred hypothetical breakpoints from co-conversions in *Exo1^-/-^*spermatocytes. The mean of each CO breakpoint distribution is indicated with a dotted line. The hotspot center is indicated with black dashed line and plotted at 0kb. Bottom, CO breakpoint maps for *Exo1^het^*, *Exo1^-/-^*, and inferred hypothetical breakpoints using co-conversions from *Exo1^-/-^* spermatocytes. The inferred gene conversion tract length based upon hypothetical CO breakpoints from *Exo1^-/-^* co-conversions (127bp) was much shorter than the directly measured co-conversion tract length of 692 ± 567bp. This shorter estimate was like the apparent estimated CO gene conversion tract length of *Exo1^-/-^* COs (182bp). This suggests that reciprocal CO asymmetry does not estimate gene conversion tract length for COs in *Exo1^-/-^*spermatocytes owing to the higher breakpoint frequency near the hotspot center. Taken together, reciprocal CO asymmetry only accurately estimates gene conversion tract length when CO breakpoint distributions are evenly distributed away from the hotspot center, such as is found when MutLgamma is proficient. CentiMorgans/Megabase (cM/Mb).

**Figure S3. Representative co-conversions isolated from all mutants**

Representative co-conversion plots isolated from the indicated genotypes and meiotic prophase stage. All mapped recombinants are included in the Supplementary Data File. The blue line plots the minimal and the gray line the maximal possible gene conversion tract. The x-axis shows ticks marking polymorphisms at the top and relative position within the hotspot at the bottom. The center of the hotspot (PRDM9 binding site polymorphism) is marked by a vertical dotted line and plotted at 0kb. Total frequency ± SD (top left) and number isolated (n_Tot_, top right) of co-conversions are shown. The number of co-conversions plotted is proportional to their frequency for visual comparison between genotypes. Diplotene samples were used wherever possible given their higher purity than Late 4C in general. Co-conversions are short and rare in WT or mutants that lack CO designation (RNF212), and common in CO maturation deficient mutants.

## Methods

### Mice

Experimental animals, C57BL/6J x DBA/2J (BxD) F1 hybrid male mice were generated by crossing inbred mouse strains C57BL/6J (B) and DBA/2J (D). Single mutants (*Exo1^-/-^*, *Hfm1^-/-^*, *Hei10^-/-^*, *Rnf212^-/-^*, *Mlh3^-/-^*, *Mlh3^DN/DN^*) were generated from independently maintained heterozygous mice in the B and D strains that were genotyped to verify homozygosity of the corresponding haplotype for the *59.5* hotspot. Double mutants, (*Mlh3^-/-^ Exo1^-/-^*, *Mus81^-/-^ Exo1^-/-^*, *Msh2^-/-^ Exo1^-/-^*) were generated similarly from mice independently maintained as heterozygous for both genes in both strains. *Mlh3^-/DN^* were generated by crossing *Mlh3^+/-^* with *Mlh3^+/DN^* mice from the D and B strains, respectively. *Msh2^-/-^* were generated by crossing *Msh2.1^tm2.1Rak/J^* with *Stra8-cre* and were maintained as a null line in both strains.

### Spermatocyte spreads and immunofluorescence

This protocol was performed as described (Zelazowski et al., 2017). Briefly, whole or part of decapsulated testes were digested with collagenase (2mg/ml) in 2ml of testis isolation medium (TIM) for 55’ at 32°C while shaken at 500rpm. (TIM: 104mM NaCl, 45mM KCl, 1.2mM MgSO4, 0.6mM KH2PO4, 0.1% (w/v) glucose, 6mM sodium lactate, 1mM sodium pyruvate, pH 7.3, filter sterilized). These extracted tubules were washed three times with 15ml TIM, then digested with Trypsin (0.7 mg/ml) in 2ml TIM containing DNAse I (4 μg/ml) for 15’ at 32°C/500rpm. This digestion was stopped by adding Trypsin inhibitor to a final concentration of 4 mg/ml in TIM and an additional 50μl of 0.4 μg/μl DNAse I was added to the mix. The suspension was then aspirated for 2’ with a transfer pipette and then filtered through 70-micron cell strainer to separate the cells. The cells were pelleted and washed twice in TIM containing DNAse I (1 μg/ml), and then resuspended in PBS. The amount of PBS used was proportional to the starting tissue, and typically 12-15ml was used for a whole WT testis and 3-6ml for mutant testis. The cells were then resuspended and incubated in hypotonic, 0.1M sucrose prewarmed to 37°C for 3-5’. In the meantime, a moist flat chamber was prepared and 65μl of 1% PFA (pH 9.2)/0.1% Triton X-100 laid onto glass slides. The sucrose cell suspension was then transferred to these glass slides, ∼20ul per slide. These slides remain in the closed moist chamber for 2.5 hours, then with the lid ajar for 30’, then with the lid open for an additional 30’. Slides were then washed with water containing 1:250 Photo-Flo 200 and air-dried before storage at −80°C.

For antibody staining, the slides were blocked with antibody dilution buffer (ADB) for 30’ at 37°C in a moist flat chamber. (ADB: 10% goat serum, 3% BSA, 0.05% Triton X-100, 1xPBS). Slides were then incubated with primary antibody diluted in ADB either overnight at room temperature or for 1 hr at 37°C. The slides were washed once with 1xPBS/0.4% Photo-Flo 200, then once with 1xPBS/0.4% Photo-Flo 200/0.01% Triton X-100, for 5’ each. Slides were blocked with ADB for 10’ at 37°C and then incubated with secondary antibody diluted in ADB. Upon secondary antibody treatment, slides were maintained in the dark. Slides were washed with 1xPBS/0.4% Photo-Flo 200 and then with 1xPBS/0.4% Photo-Flo 200/ 0.01% Triton X-100, for 5’ as described above. These slides were then air-dried and mounted with coverslips using Prolong Gold antifade containing DAPI.

### Metaphase spreads

This protocol was performed as previously described (Zelazowski et al., 2017). Briefly, whole or part of decapsulated testes were incubated at room temperature in a 2.9% isotonic sodium citrate solution. The tubules were pulled apart until sufficient cells were released into the solution, making it turbid. The supernatant was then transferred to a 15ml tube while avoiding tubules and pelleted. Cells were resuspended in 1% hypotonic sodium citrate solution for 12’ at room temperature. Cells were pelleted, resuspended, thoroughly mixed, and incubated in fixative for 5’ (Fixative: 3 parts 200 proof ethanol, 1 parts glacial acetic acid, 0.025 parts chloroform). After the first incubation, cells were resuspended and incubated in fresh fixative for 5’. Approximately 20μl of this suspension was dropped on a grease-free slide, air-dried, and stained with mild shaking in 1:20 dilution Giemsa in 1x PBS (pH 6.5) for 45’. Slides were then briefly washed with water, dried, and mounted with a cover slip for imaging.

### Synchronization of spermatogenesis

Mouse spermatocytes were synchronized as described (Hogarth et al., 2013; Patel et al., 2019). Briefly, WIN 18,446 was pipet fed to 1-3 days postpartum (dpp) BxD F1 male neonates at 100μg/g body weight for seven (or more, if needed for weight gain) consecutive days every 22 - 24 hours. Starting on the 8^th^ day of treatment, if the mice weighed at least 4g, they were intraperitoneally injected with 200μg of retinoic acid suspended in 10μl dimethyl sulfoxide. The duration of days post-injection that allows enrichment for specific prophase stages was determined based on published reports on spermatocyte prophase timing (Hogarth, 2013) and experimentation. Typically, synchrony is maintained for at least 6 rounds of spermatogenesis allowing isolation of adult populations. For synchronization of *Hei10^-/-^*, *Rnf212^-/-^*, *Mlh3^-/-^* and *Exo1^-/-^*, male neonates from experimental crosses were genotyped between 0-2dpp to test for the presence of animals homozygous for null allele before treatment.

### Isolation of stage-specific and Late 4C spermatocytes by flow cytometry

Spermatocytes were isolated as described in earlier reports (Cole et al., 2014; Zelazowski et al., 2017). Briefly, seminiferous tubules from decapsulated testis were digested with 0.5 mg/ml collagenase in Gey’s Balanced Salt Solution (GBSS) for 15 minutes at 33°C, while shaking at 500 RPM. For collagenase digestion and most subsequent steps, ∼10ml of GBSS was used per decapsulated WT or mutant mouse testis. These tubules were then washed once with GBSS, followed by digestion with trypsin at 0.5 mg/ml in GBSS supplemented with 1 μl/ml DNase I, for 15 minutes at 33°C/500 RPM. Subsequently, Newborn Calf Serum (NCS) is added to a concentration of 5% to inactivate Trypsin. To individualize cells, the solution was pipetted up and down for 3’ with a transfer pipette and then filtered through a 70-micron cell strainer. Cells were then washed multiple times in GBSS with 5% NCS/1 μg/ml DNAse I and subsequently stained by incubating in a solution of 5 μg/ml Hoechst 33342 in GBSS with 5% NCS/1 μg/ml DNAse I for 45’ minutes at 33°C/500 RPM. The amount of Hoechst staining solution was proportional to tissue weight, 6ml and 3ml were typically used for staining one WT or mutant testis, respectively. After Hoechst staining, propidium iodide was added to a final concentration of 0.2 μg/ml and the cells were filtered through a 70-micron cell strainer. Cells were then sorted in a BD Aria or BD Fusion flow cytometer, with UV 350nm argon laser. On a linear scale, blue (DNA content) and red (chromosome compaction) emission spectra were used to gate for different stages of meiosis, with for example, leptonema appearing closest to the left and diplonema the right of the red emission spectrum. Isolation of synchronized mouse spermatocytes produces the highest purity of specific stages; however the gating strategy can successfully enrich for late stages (pachynema and diplonema) albeit with a slight loss in purity by comparison. Empirical optimization by combining the sort-gating strategy and subsequent purity analysis is required to optimize and verify sample enrichment. Sorted cells were washed with PBS, pelleted at 600g for 5’, and snap-frozen as a dry pellet in dry ice/ethanol slurry for storage at −80°C. A portion, typically 5,000-20,000 cells were surface spread and stained with antibodies (SYCP3/SYCP1/Histone H1t/gammaH2AX) and DAPI to test for the purity of the sorted population.

### Genomic DNA isolation

The genomic DNA was isolated as previously described (Zelazowski et al., 2017). Cell pellets were resuspended in 500μl 0.2x SSC, pH 7 (SSC 20x: 3M NaCl and 0.3M citric acid trisodium salt dihydrate). For this suspension, 60μL β-mercaptoethanol, 10μL of 20 mg/ml Proteinase K and 50μL of 10% SDS were added sequentially and incubated for 30’ at 55°C/500rpm. This cell digest was then extracted with Phenol/chloroform/isoamyl alcohol. DNA was then precipitated with two volumes of ice-cold 100% ethanol with linear polyacrylamide as a DNA carrier. The DNA pellet was washed with 70% ethanol, air-dried for ∼5’, and resuspended overnight in 5mM Tris (pH 7.5) at 4°C.

### Determining amplification efficiency of DNA

Amplification efficiency on isolated DNA was performed as described (Cole and Jasin, 2011). Briefly, using nanodrop estimate of DNA concentration as a starting point and 12pg DNA (2 diploid genome equivalents) as input per PCR reaction, 24-48 PCR reactions were performed. The amplification consisted of 2 rounds of nested PCRs, with conditions identical to those used for NCO/CO amplification. Each round of this PCR consisted of one allele-specific primer (ASP) and one universal primer, and hence amplification efficiency estimated this way applies only to these primary and secondary ASPs. The amplified products were run on a gel and the number of negative wells were counted. The amplification adjustment factor is then given by (-log_2_(Number of negative wells/Number of total wells))/2. For maximum reproducibility, amplification adjustment factor must fall between 0.2 and 0.8, if not amplifiable DNA concentration is revised and the amplification efficiency test is repeated.

### CO assay

CO assay was performed as described (Cole and Jasin, 2011; Cole et al., 2010). Sperm and spermatocyte DNA was analyzed in sufficiently small pools such that the number of wells with more than one crossover was kept to a minimum. As such, input DNA was higher in CO defective mutants like *Exo1^-/-^* (∼100 genomic DNA equivalents) and *Mlh3^-/-^* (∼300) as compared to *Exo1^nd/nd^* (∼40). The primer annealing temperature was empirically determined for each batch of 11.1x buffer. For each primary PCR reaction (8μl) DNA was added to 1x buffer (10x: 450mM Tris-HCl pH 8.8, 110mM (NH4)2SO4, 45mM MgCl2, 67mM Beta-mercaptoethanol, 44μM EDTA, 10mM each: dATP, dTTP, dGTP, and dCTP, and 1.13 mg/ml non-acetylated BSA), 12.5mM Tris-base, 0.2μM of each primer, 0.25U of Taq, and 0.05U of Pfu polymerase. 2μl of the primary PCR reaction was digested with S1 nuclease (10μl reaction): 0.7U/μl S1 nuclease in 1x buffer: 20mM sodium acetate, 1mM Zn acetate, 0.1M NaCl, and the reaction was diluted and stopped with 45μL of dilution buffer (10mM Tris-HCl pH 7.5 and 5 μg/ml sonicated salmon sperm DNA). 1.6μl of the diluted primary PCR was used as input for secondary PCR using identical conditions as described above but with nested ASPs. The secondary PCR plate wells were tested for amplification, by running a 1/10 of the reaction on a 1% agarose gel. Tertiary PCR (30μl) was performed on positive secondary PCR wells. The PCR program for primary, secondary, and tertiary PCRs were denaturation (96°C, 1’ for the first denaturation and 20’’ for subsequent steps), annealing (30’’ at optimized temperature), extension (65°C, 1’ per kb). Both primary and secondary PCRs had 27 cycles of amplification, while the tertiary PCR had 30 cycles of amplification. The tertiary PCR plate, in its 96 well format is then dot blotted to positively charged nylon membrane to genotype by southern blotting using allele-specific oligos (ASO) probes. An equal amount of somatic DNA was used as the negative control.

### NCO assay

NCO assay was done as described (Cole and Jasin, 2011; Cole et al., 2010). NCOs were amplified much like the COs described above, except for a few key differences: 1) The primer pairs used have one ASP and one universal primer for unbiased amplification of COs, NCOs, and non-recombinant DNA; 2) The input DNA was no higher than 30 haploid genome equivalents per well, so the recombinant signal was detectable over the non-recombinant background; 3) NCO assays include only a primary (8μl) and secondary (30μl) PCRs without intervening S1 nuclease digestion, with 27 and 36 cycles of amplification, respectively; and 4) The entire secondary PCR was dot blotted and genotyped by Southern blotting using ASO probes.

### Genotyping with ASO probes

Genotyping amplicons from CO/NCO assays were done as described (Cole and Jasin, 2011; Cole et al., 2010). The ASOs were radiolabeled with T4 polynucleotide kinase in a 10μl reaction (Buffer: 70mM Tris-HCl, pH 7.5, 10mM MgCl2, 5mM spermidine trichloride, 2mM dithiothreitol; 8ng ASO; 0.35μL T4 polynucleotide kinase (10U/μl), 0.2μL γ-32P-ATP (10mCi/ml)) per two blots. Reaction was done at 37°C for 45-60’ and was stopped with 20μl stop buffer (25mM EDTA, 0.1% SDS, 10μM ATP) and then 20μL of unlabeled ASO of the opposite genotype (8 μg/ml) was added to the mix.

Dot blot membranes were rinsed in 2X SSC (20x: 3M NaCl and 0.3M citric acid trisodium salt dihydrate) and pre-hybridized for 15’ with Tetramethylammonium Chloride (TMAC) hybridization buffer (3M TMAC, 0.6% SDS, 10mM NaPO4 pH 6.8, 1mM EDTA, 4 μg/ml yeast RNA, in 5x Denhardt’s solution) at 56°C for 10–15’. 50x Denhardt’s solution: 1% (w/v) Ficoll 400, 1% (w/v) polyvinyl pyrrolidone, 1% (w/v) BSA (Fraction V); filter sterilized. Next, the buffer was replaced with fresh TMAC hybridization buffer with sonicated salmon sperm DNA at a concentration of 8.4 μg/ml, and incubated for 10’ at 53°C. Radio-labeled ASOs were then added, and incubated for 45-60’ at 53°C. Next, the blots were washed thrice with TMAC wash buffer (3M TMAC, 0.6% SDS, 10mM NaPO4 pH 6.8, 1mM EDTA), within a span of ∼20’ while incubating the blots at 56°C. For the final wash, the blots were incubated with TMAC wash buffer for 15’ at 56°C. Blots were rinsed twice with 2X SSC, excess liquid was blotted away, wrapped in plastic and exposed overnight on a phosphorimager screen.

### Cytological analysis

Immunofluorescence experiments were performed blinded. All animals were BxD. Mutants and controls within an experimental set were analyzed by one individual to reduce inter-experimentalist variation. Only on axis (SYCP3-associated) foci were counted and any X/Y MLH1 or MLH3 foci were excluded from the analysis.

For DMC1 foci, SYCP3 staining was used to stage cells and zygonema cells were chosen such that some chromosomes have contiguous SYCP3 axis and were categorized as early-mid zygonema (<50% synapsis, visually estimated) or mid-late zygonema (>50% synapsis, visually estimated). Similar numbers (Fishers Exact test, two-tailed, ns) of each zygotene stage were compared between WT (38 and 39, respectively), *Exo1^het^* (16 and 24 respectively), and *Exo1^-/-^* (33 and 40, respectively).

For MLH1 and MLH3 foci, every cell that was positive for foci were counted.

Metaphase bivalents counts were performed blinded as much as possible by comparing together mutants with similar severity of CO loss (for example, *Mlh3^DN/DN^* and *Exo1^-/-^Mus81^-/-^*). To minimize inter-experimentalist variation, the authors agreed upon a common strategy for counting. To ensure we only analyzed intact nuclei, independent chromosomes structures were quantified first as ‘objects’ and subsequently were categorized as either bivalents or univalents. A cell was then included only if the number of total bivalents and univalents accounted for all chromatids expected (40 for mouse spermatocytes). Cells that had chromosome structures that the author was unable to classify as a bivalent or univalent were not included in the analysis.

Co-efficient of Coincidence (CoC) analysis was done as described (White et al., 2017). Briefly, the DAPI rich end of each chromosome (centromere) was designated the starting point and the distance of each focus from the start and length of the chromosome axis was quantified manually using ImageJ. Based on the number of MLH1 foci per micron axis and cells counted (61 and 42 cells for WT and *Exo1^-/-^*, respectively), 10 intervals per bivalent were assessed. The Mean CoC vs inter-interval distance was calculated using MATLAB (White et al., 2017). Axis lengths for WT and *Exo1^-/-^* spermatocytes (**Figure S1A**) were acquired from the CoC dataset.

### Experimental design, quantification, and statistical analysis

Cytological analyses were performed blinded, and samples were cross checked by independent experimentalists. For recombination analysis, NCO and CO plates from samples of different mutants were amplified, hybridized, and genotyped by Southern blotting together to reduce inter-experimental variation. While the genotyping of recombinants wasn’t blinded, most mutants had similar severity of CO loss and different phenotypes weren’t apparent until results were fully tabulated.

All data are shown as means with standard deviation and 95% confidence intervals. The p-value is shown when the difference was significant and are adjusted for multiple comparisons as appropriate. Broadly, we used Fisher’s exact test, two-tailed for categorical comparisons. To compare means, we used Mann-Whitney for pairwise and Kruskal-Wallis followed by Dunn’s multiple comparison correction for non-pairwise comparisons. We have reported the tests performed in text and figure legends. We show the number of animals analyzed as capital ‘N’, and number of quantified data points as small ‘n’. For most conditions, at least three animals were analyzed per genotype and all animals were BxD.

## Acknowledgments

We thank P. Cohen for mice and the MLH3 antibody, W. Edelmann, N. Hunter, and R. Pezza for mice, M.A. Handel for the H1t antibody, and members of the Cole lab for comments on the manuscript. We also thank N. Kleckner, N. Hunter, G.V. Börner, J. Ahuja, and A. Villeneuve for sharing their insights. This work was supported by National Institutes of Health grants R01HD098129 and DP2HD087943 (F.C.) and summer undergraduate fellowship R25CA181004 (H.D.), the Cockrell Endowment (T.P.) and H.E.B Fellowships (T.P. and M.F.), CPRIT grant R1213 (F.C.) and training grant RP170067 (R.K., V.K., I.Y.). We acknowledge the National Institutes of Health CA16672 for support of the Research Animal Support Facility Smithville and the CPRIT RP170628 for support of the Flow Cytometry and Cellular Imaging Core.

## Author contributions

T.P., L.P., R.K., E.H., H.D., I.F., I.O., H.N., M.F., and P.C. conducted experiments; T.P., and F.C. performed statistical analyses; and T.P., L.P., R.K., and F.C. designed the experiments and analyzed the data. T.P. and F.C. wrote the paper.

---

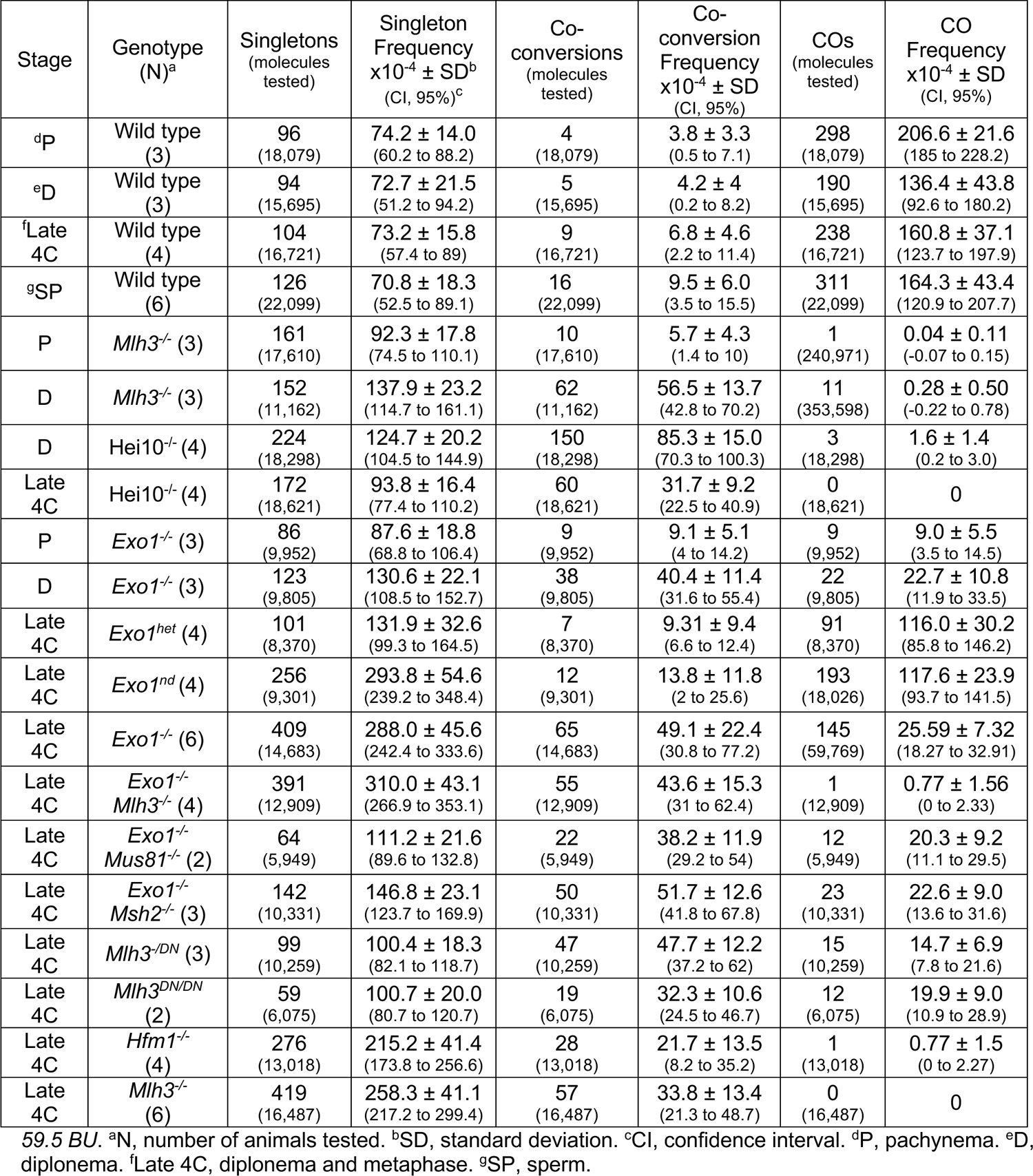

