## Supplement figures and tables for "Genetic dissection of crossover mutants defines discrete intermediates in mouse meiosis"

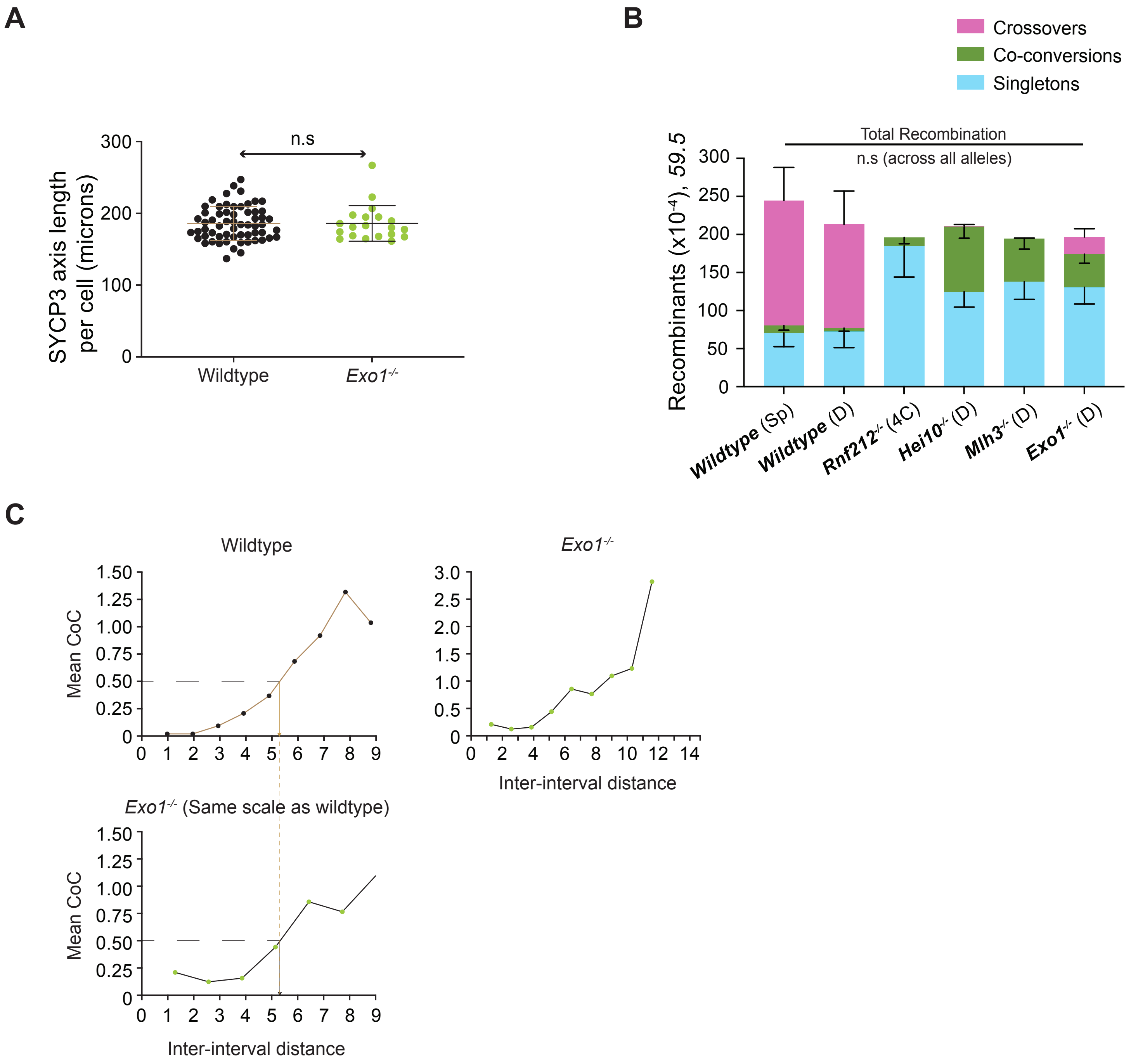

Figure S1 Premkumar et al

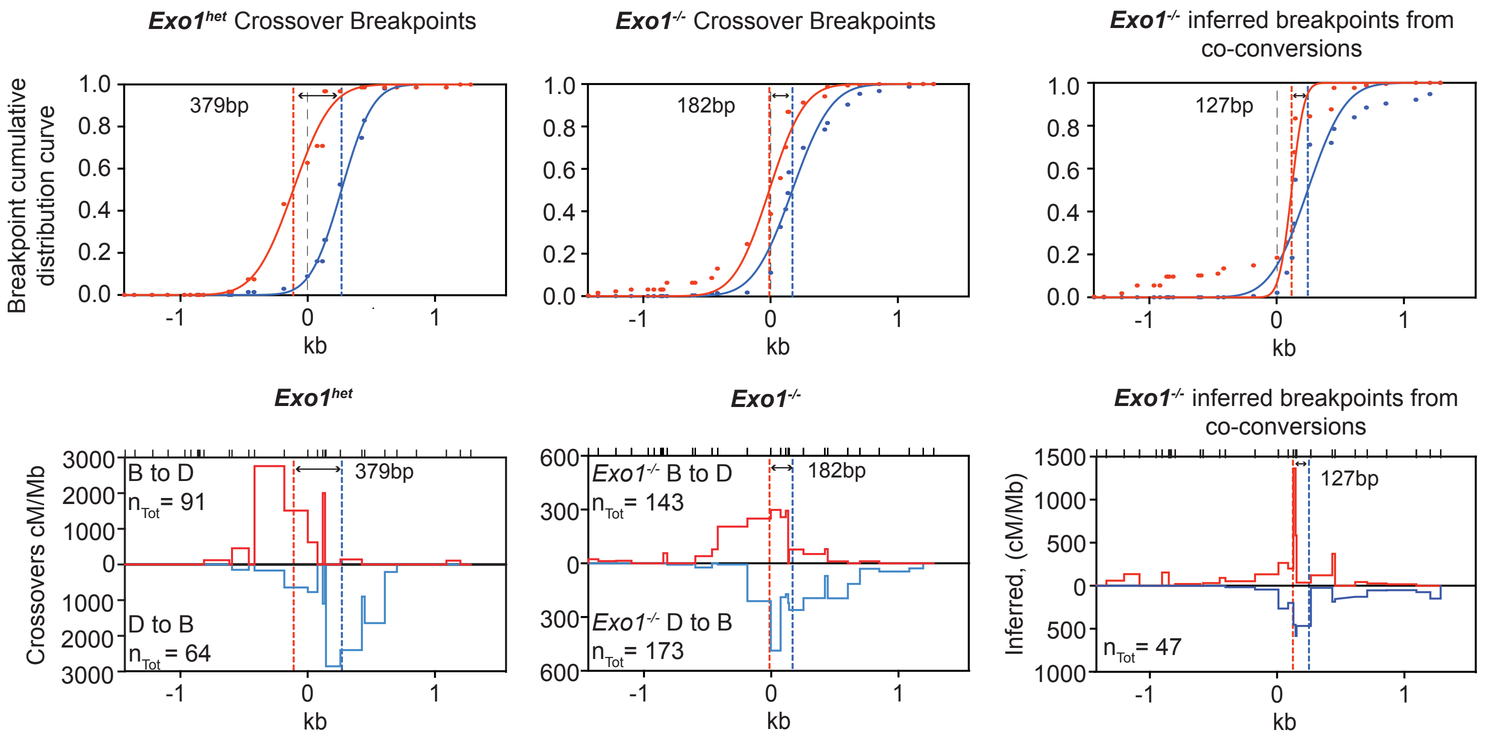

Figure S2 Premkumar et al

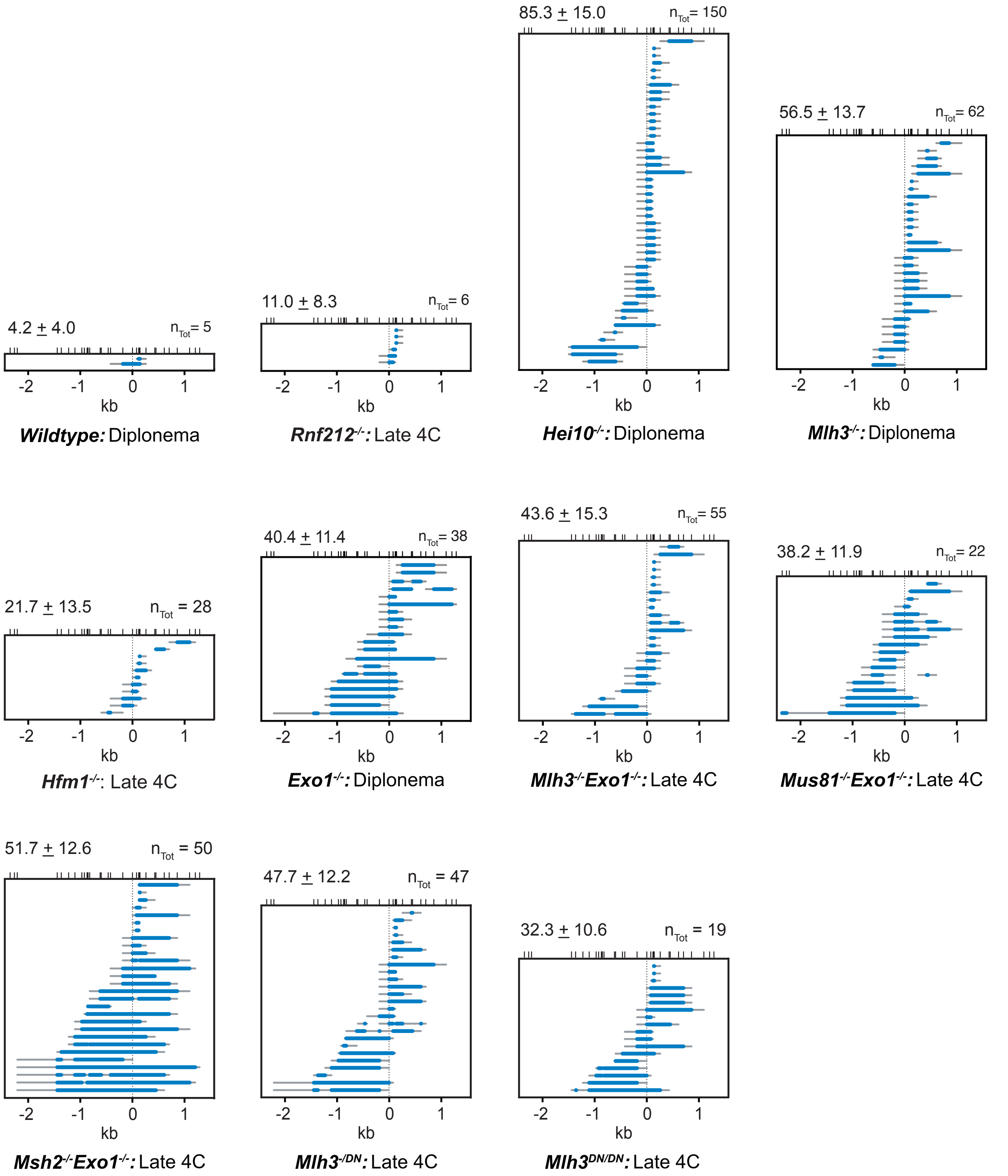

Figure S3 Premkumar et al

| Wildtype pachynema post sort (raw count) | | | | | | | | |
|---|---|---|---|---|---|---|---|---|
| Sample | Target stage | Pre-Leptonema | Leptonema | Zygonema | Pachynema | Diplonema | Metaphase | 2º spermatocytes & spermatids |
| #1 28 dpi | Pachynema | 0 | 0 | 5 | **66** | 9* | 0 | 8 |
| #2 30 dpi | Pachynema | 0 | 0 | 1 | **78** | 24* | 0 | 0 |
| #3 37 dpi | Pachynema | 0 | 0 | 3 | **162** | 38* | 0 | 0 |
| Purity | 76.8±2.6%* | | | | | | | |
| Pre-sort (raw count) | | | | | | | | |
| #1 28 dpi | Pachynema | 0 | 0 | 4 | **156** | 0 | 0 | 0 |
| #2 30 dpi | Pachynema | ND | ND | ND | ND | ND | ND | ND |
| #3 37 dpi | Pachynema | 0 | 0 | 7 | **93** | 0 | 0 | 0 |
| Purity | 95.3±3.2% | | | | | | | |

| Wildtype diplonema post sort (raw count) | | | | | | | | |
|---|---|---|---|---|---|---|---|---|
| Sample | Target stage | Pre-Leptonema | Leptonema | Zygonema | Pachynema | Diplonema | Metaphase | 2º spermatocytes & spermatids |
| #1 43 dpi | Diplonema | 0 | 0 | 0 | 1 | **98** | 14 | 6 |
| #2 17 dpi | Diplonema | 0 | 0 | 7 | 0 | **64** | 5 | 0 |
| #3 33 dpi | Diplonema | 0 | 0 | 0 | 0 | **82** | 21 | 2 |
| Purity | 81.6±3.1% | | | | | | | |
| Pre-sort (raw count) | | | | | | | | |
| #1 43 dpi | Diplonema | ND | ND | ND | ND | **ND** | ND | ND |
| #2 17 dpi | Diplonema | 0 | 4 | 96 | 13 | **36** | 6 | 5 |
| #3 33 dpi | Diplonema | 0 | 0 | 97 | 0 | **60** | 3 | 0 |
| Purity | 30±10.6% | | | | | | | |

| Wildtype Late 4C (diplonema & metaphase) post sort (raw count) | | | | | | | | |
|---|---|---|---|---|---|---|---|---|
| Sample | Target stage | Pre-Leptonema | Leptonema | Zygonema | Pachynema | Diplonema | Metaphase | 2º spermatocytes & spermatids |
| H | Late-4C | 0 | 0 | 2 | 15 | **80** | **3** | 0 |
| I | Late-4C | 0 | 0 | 3 | 15 | **67** | **3** | 0 |
| J | Late-4C | 0 | 0 | 7 | 0 | **43** | **1** | 0 |
| K | Late-4C | ND | ND | ND | ND | ND | ND | ND |
| Purity | 82.9±3.3% | | | | | | | |

| *Mlh3*$^{-/-}$ pachynema post sort (raw count) | | | | | | | | |
|---|---|---|---|---|---|---|---|---|
| Sample | Target stage | Pre-Leptonema | Leptonema | Zygonema | Pachynema | Diplonema | Metaphase | 2º spermatocytes & spermatids |
| #1 29 dpi | Pachynema | 0 | 4 | 0 | **80** | 16* | 0 | 0 |
| #2 20 dpi | Pachynema | 0 | 4 | 0 | **59** | 37* | 0 | 0 |
| #3 46 dpi | Pachynema | 0 | 4 | 0 | **32** | 64* | 0 | 0 |
| Purity | 57.0±24.1%* | | | | | | | |
| Pre-sort (raw count) | | | | | | | | |
| #1 29 dpi | Pachynema | 0 | 0 | 0 | **163** | 0 | 0 | 0 |
| #2 20 dpi | Pachynema | 0 | 0 | 0 | **159** | 2 | 0 | 0 |
| #3 46 dpi | Pachynema | 0 | 2 | 0 | **160** | 0 | 0 | 0 |
| Purity | 99.2±0.7% | | | | | | | |

| *Mlh3*$^{-/-}$ diplonema post sort (raw count) | | | | | | | | |
|---|---|---|---|---|---|---|---|---|
| Sample | Target stage | Pre-Leptonema | Leptonema | Zygonema | Pachynema | Diplonema | Metaphase | 2° spermatocytes & spermatids |
| #1 23 dpi | Diplonema | 0 | 0 | 0 | 0 | **100** | 0 | 0 |
| #2 40 dpi | Diplonema | 0 | 0 | 0 | 0 | **98** | 2 | 0 |
| #3 49 dpi | Diplonema | 0 | 0 | 0 | 0 | **97** | 3 | 0 |
| Purity | 98.3±1.5% | | | | | | | |
| Pre-sort (raw count) | | | | | | | | |
| #1 23 dpi | Diplonema | 0 | 1 | 0 | 22 | **139** | 0 | 0 |
| #2 40 dpi | Diplonema | 0 | 2 | 56 | 2 | **100** | 0 | 0 |
| #3 49 dpi | Diplonema | 0 | 10 | 62 | 0 | **39** | 0 | 0 |
| Purity | 61.1±25.3% | | | | | | | |

| *Mlh3*$^{-/-}$ Late 4C (diplonema & metaphase) post sort (raw count) | | | | | | | | |
|---|---|---|---|---|---|---|---|---|
| Sample | Target stage | Pre-Leptonema | Leptonema | Zygonema | Pachynema | Diplonema | Metaphase | 2° spermatocytes & spermatids |
| ID 3125 | Late-4C | 0 | 0 | 0 | 0 | **38** | **2** | 0 |
| ID 3126 | Late-4C | 0 | 0 | 0 | 0 | **43** | **2** | 0 |
| ID 3123 | Late-4C | 0 | 0 | 0 | 0 | **38** | **6** | 0 |
| ID 159 | Late-4C | 0 | 0 | 0 | 0 | **39** | **1** | 0 |
| ID 189 | Late-4C | 0 | 0 | 0 | 1 | **39** | **0** | 0 |
| ID 842 | Late-4C | 0 | 0 | 0 | 1 | **49** | **2** | 0 |
| Purity | 99.3±1.1% | | | | | | | |

| *Exo1*$^{-/-}$ pachynema post sort (raw count) | | | | | | | | |
|---|---|---|---|---|---|---|---|---|
| Sample | Target stage | Pre-Leptonema | Leptonema | Zygonema | Pachynema | Diplonema | Metaphase | 2° spermatocytes & spermatids |
| #1 37 dpi | Pachynema | 0 | 0 | 0 | **46** | 0 | 0 | 0 |
| #2 37 dpi | Pachynema | 0 | 0 | 0 | **40** | 0 | 0 | 0 |
| #3 37 dpi | Pachynema | 0 | 0 | 0 | **48** | 0 | 0 | 0 |
| Purity | 100.0±0.0% | | | | | | | |
| Pre-sort (raw count) | | | | | | | | |
| #1 37 dpi | Pachynema | 0 | 0 | 0 | **71** | 2 | 0 | 0 |
| #2 37 dpi | Pachynema | 0 | 0 | 2 | **39** | 0 | 0 | 0 |
| #3 37 dpi | Pachynema | 0 | 0 | 0 | **56** | 0 | 1 | 0 |
| Purity | 96.9±1.6% | | | | | | | |

| *Exo1*$^{-/-}$ diplonema post sort (raw count) | | | | | | | | |
|---|---|---|---|---|---|---|---|---|
| Sample | Target stage | Pre-Leptonema | Leptonema | Zygonema | Pachynema | Diplonema | Metaphase | 2° spermatocytes & spermatids |
| #1 41 dpi | Diplonema | 0 | 0 | 0 | 0 | **66** | 4 | 0 |
| #2 41 dpi | Diplonema | 0 | 0 | 0 | 0 | **42** | 0 | 0 |
| #3 41 dpi | Diplonema | 0 | 0 | 0 | 0 | **47** | 1 | 0 |
| Purity | 97.4±2.9% | | | | | | | |
| Pre-sort (raw count) | | | | | | | | |
| #1 41 dpi | Diplonema | 0 | 0 | 13 | 0 | **36** | 1 | 0 |
| #2 41 dpi | Diplonema | 0 | 0 | 17 | 0 | **32** | 0 | 0 |
| #3 41 dpi | Diplonema | 0 | 2 | 16 | 0 | **38** | 2 | 0 |
| Purity | 67.6±3.8% | | | | | | | |

| *Exo1*$^{-/-}$ Late 4C (diplonema & metaphase) post sort (raw count) |
|---|

| Sample | Target stage | Pre-Leptonema | Leptonema | Zygonema | Pachynema | Diplonema | Metaphase | 2º spermatocytes & spermatids |
|---|---|---|---|---|---|---|---|---|
| ID 2501 | Late-4C | 0 | 0 | 0 | 0 | **40** | **3** | 0 |
| ID 2325 | Late-4C | 0 | 0 | 0 | 6 | **50** | **2** | 0 |
| ID 2263 | Late-4C | 0 | 0 | 1 | 12 | **45** | **5** | 0 |
| ID 2089 | Late-4C | 0 | 0 | 0 | 0 | **50** | **3** | 0 |
| ID 2088 | Late-4C | 0 | 0 | 0 | 3 | **50** | **5** | 0 |
| ID 2104 | Late-4C | 0 | 0 | 0 | 4 | **54** | **2** | 0 |
| ID 2105 | Late-4C | 0 | 0 | 0 | 0 | **56** | **1** | 0 |
| ID 2078 | Late-4C | 0 | 0 | 0 | 0 | **47** | **3** | 0 |
| Purity | 94.6±7.3% | | | | | | | |

| *Exo1^het^* Late 4C (diplonema & metaphase) post sort (raw count) | | | | | | | | |
|---|---|---|---|---|---|---|---|---|
| Sample | Target stage | Pre-Leptonema | Leptonema | Zygonema | Pachynema | Diplonema | Metaphase | 2º spermatocytes & spermatids |
| ID 2077 | Late-4C | 0 | 0 | 0 | 5 | **45** | **2** | 0 |
| ID 2087 | Late-4C | 0 | 0 | 0 | 2 | **37** | **2** | 0 |
| ID 2437 | Late-4C | 0 | 3 | 0 | 2 | **36** | **0** | 0 |
| ID 2319 | Late-4C | 0 | 1 | 0 | 0 | **44** | **1** | 0 |
| Purity | 92.8±4.5% | | | | | | | |

| *Mlh3^-/-^Exo1^-/-^* Late 4C (diplonema & metaphase) post sort (raw count) | | | | | | | | |
|---|---|---|---|---|---|---|---|---|
| Sample | Target stage | Pre-Leptonema | Leptonema | Zygonema | Pachynema | Diplonema | Metaphase | 2º spermatocytes & spermatids |
| ID 2620 | Late-4C | 0 | 0 | 0 | 1 | **42** | **4** | 0 |
| ID 2801 | Late-4C | 0 | 0 | 0 | 1 | **52** | **2** | 0 |
| ID 3400 | Late-4C | 0 | 0 | 1 | 0 | **60** | **1** | 0 |
| ID 3298 | Late-4C | 0 | 0 | 0 | 0 | **45** | **3** | 0 |
| Purity | 98.6±0.9% | | | | | | | |

| *Mus81^-/-^Exo1^-/-^* Late 4C (diplonema & metaphase) post sort (raw count) | | | | | | | | |
|---|---|---|---|---|---|---|---|---|
| Sample | Target stage | Pre-Leptonema | Leptonema | Zygonema | Pachynema | Diplonema | Metaphase | 2º spermatocytes & spermatids |
| ID 442 | Late-4C | 0 | 0 | 0 | 0 | **40** | **5** | 0 |
| ID 1660 | Late-4C | 0 | 0 | 4 | 8 | **36** | **3** | 0 |
| Purity | 88.2±16.6% | | | | | | | |

| *Msh2^-/-^Exo1^-/-^* Late 4C (diplonema & metaphase) post sort (raw count) | | | | | | | | |
|---|---|---|---|---|---|---|---|---|
| Sample | Target stage | Pre-Leptonema | Leptonema | Zygonema | Pachynema | Diplonema | Metaphase | 2º spermatocytes & spermatids |
| ID 2488 | Late-4C | 0 | 0 | 0 | 0 | **40** | **2** | 0 |
| ID 2489 | Late-4C | 0 | 0 | 0 | 1 | **40** | **0** | 0 |
| ID 2290 | Late-4C | 0 | 0 | 1 | 0 | **40** | **0** | 0 |
| Purity | 98.3±1.4% | | | | | | | |

| *Mlh3^-/DN^* Late 4C (diplonema & metaphase) post sort (raw count) | | | | | | | | |
|---|---|---|---|---|---|---|---|---|
| Sample | Target stage | Pre-Leptonema | Leptonema | Zygonema | Pachynema | Diplonema | Metaphase | 2º spermatocytes & spermatids |
| ID 32 | Late-4C | 0 | 0 | 0 | 1 | **37** | **3** | 0 |
| ID 88 | Late-4C | 0 | 0 | 0 | 1 | **39** | **3** | 0 |

| | | | | | | | | |
|---|---|---|---|---|---|---|---|---|
| ID 90 | Late-4C | 0 | 0 | 0 | 1 | **38** | **1** | 0 |
| Purity | 97.6±0.08% | | | | | | | |

| *Mlh3*$^{DN/DN}$ Late 4C (diplonema & metaphase) post sort (raw count) | | | | | | | | |
|---|---|---|---|---|---|---|---|---|
| Sample | Target stage | Pre-Leptonema | Leptonema | Zygonema | Pachynema | Diplonema | Metaphase | 2º spermatocytes & spermatids |
| ID 445 | Late-4C | 0 | 0 | 0 | 0 | **43** | **1** | 0 |
| ID 409 | Late-4C | 0 | 0 | 0 | 0 | **35** | **7** | 0 |
| Purity | 100.0±0.0% | | | | | | | |

| *Hfm1*$^{-/-}$ Late 4C (diplonema & metaphase) post sort (raw count) | | | | | | | | |
|---|---|---|---|---|---|---|---|---|
| Sample | Target stage | Pre-Leptonema | Leptonema | Zygonema | Pachynema | Diplonema | Metaphase | 2º spermatocytes & spermatids |
| ID 441 | Late-4C | 0 | 0 | 0 | 0 | **43** | **1** | 0 |
| ID 443 | Late-4C | ND | ND | ND | ND | ND | ND | ND |
| ID 514 | Late-4C | 0 | 0 | 0 | 0 | **38** | **3** | 0 |
| ID 504 | Late-4C | 0 | 0 | 0 | 0 | **43** | **6** | 0 |
| Purity | 100.0±0.0% | | | | | | | |

| *Exo1*$^{nd/nd}$ Late 4C (diplonema & metaphase) post sort (raw count) | | | | | | | | |
|---|---|---|---|---|---|---|---|---|
| Sample | Target stage | Pre-Leptonema | Leptonema | Zygonema | Pachynema | Diplonema | Metaphase | 2º spermatocytes & spermatids |
| ID 161 | Late-4C | 0 | 0 | 0 | 0 | **36** | **5** | 0 |
| ID 120 | Late-4C | 0 | 0 | 0 | 1 | **34** | **5** | 0 |
| ID 254 | Late-4C | 0 | 0 | 0 | 2 | **40** | **2** | 0 |
| ID 301 | Late-4C | 0 | 0 | 0 | 0 | **42** | **0** | 0 |
| Purity | 100.0±0.0% | | | | | | | |

| *Hei10*$^{-/-}$ Late 4C (diplonema & metaphase) post sort (raw count) | | | | | | | | |
|---|---|---|---|---|---|---|---|---|
| Sample | Target stage | Pre-Leptonema | Leptonema | Zygonema | Pachynema | Diplonema | Metaphase | 2º spermatocytes & spermatids |
| KO A | Late-4C | 0 | 0 | 7 | 0 | **34** | **4** | 0 |
| KO B | Late-4C | 0 | 0 | 10 | 0 | **37** | **1** | 0 |
| KO C | Late-4C | 0 | 0 | 9 | 21 | **13** | **1** | 0 |
| KO D | Late-4C | 0 | 0 | 6 | 0 | **39** | **1** | 0 |
| Purity | 70.6±26.1% | | | | | | | |

| *Hei10*$^{-/-}$ diplonema post sort (raw count) | | | | | | | | |
|---|---|---|---|---|---|---|---|---|
| Sample | Target stage | Pre-Leptonema | Leptonema | Zygonema | Pachynema | Diplonema | Metaphase | 2º spermatocytes & spermatids |
| ID 92 | Late-4C | 0 | 0 | 0 | 0 | **92** | 14 | 0 |
| ID 94 | Late-4C | 0 | 0 | 0 | 1 | **71** | 31 | 0 |
| ID 198 | Late-4C | 0 | 0 | 0 | 1 | **97** | 0 | 0 |
| ID 462 | Late-4C | 0 | 0 | 6 | 1 | **76** | 19 | 0 |
| Purity | 82.3±13.3% | | | | | | | |
| Pre-sort (raw count) | | | | | | | | |
| Sample | Target stage | Pre-Leptonema | Leptonema | Zygonema | Pachynema | Diplonema | Metaphase | 2º spermatocytes & spermatids |

| | | | | | | | | |
|---|---|---|---|---|---|---|---|---|
| ID 92 | Late-4C | ND | ND | ND | ND | ND | ND | ND |
| ID 94 | Late-4C | 0 | 1 | 50 | 12 | **48** | 10 | 0 |
| ID 198 | Late-4C | ND | ND | ND | ND | ND | ND | ND |
| ID 462 | Late-4C | 0 | 0 | 21 | 0 | **62** | 14 | 0 |
| Purity | 51.8±17.1% | | | | | | | |

*In our original chromosome spread analysis, post-sort pachytene spermatocytes occasionally show slight splaying of the axis termini. These were scored as diplonema, however, based upon pre-sort purity, the co-existing populations within the testis (e.g., zygonema from the subsequent wave of spermatogenesis are always found in conjunction with diplonema), and the disparate distribution of recombination outcomes (**Figure 4B, 4C**) these cells are likely pachynema. In subsequent experiments, we verified that these splayed ends retain SYCP1 staining.

**Table S2.**
Allele-specific primers (ASPs) used to amplify recombinants at *59.5*

| ASP | Primer sequence (5'->3') | 5' location (GRCm39) | EVA RefSNP release 3 |
|---|---|---|---|
| Bf14590.1 | TGTTTCTGAAGCACGGGA | 19:59428515 | rs50050489 |
| Caf14590.1 | TGTTTCTGAAGCACGGGG | | |
| Bf14913.1 | CAAGACCCGGTCAGAACC | 19:59428838 | rs51436899 |
| Caf14913.1 | CAAGACCCGGTCAGAACA | | |
| Br 19630.1 | CTGGCTGACTCCATAAAGA | 19:59433590 | rs37215264 |
| Car19630.1 | CTGGCTGACTCCATAAAGG | | |
| Br19683 | GCACTGGGGATGTAATAGG | 19:59433643 | rs51754290 |
| Car19683 | GCACTGGGGATGTAATAGT | | |
| **Universal Primers** | | | |
| 59.5Uf15721 | CTGTGTACTATCATTCCTGGC | 19:59429643 | |
| 59.5Uf16055 | TGGGACTCACATGGTAAAGTG | 19:59429977 | |
| 59.5Ur18935 | CAACGAGAACACATCTGTGCCC | 19:59432898 | |
| 59.5Ur19001 | CCGCTGTGAACTGGGCGC | 19:59432960 | |

Genotyping primers used to test genotype alleles

| Allele | Primer name, sequence (5'->3') | $T_m$ | Bands | Multiplex |
|---|---|---|---|---|
| HFM1$^{Gt(OST347241)Lex}$ | F1: GCTGTCCAGTACTTTTATACAAC | 60°C, 35 cycles | WT 340bp, MT 270bp | Yes |
| | R1: GGTACAAGCTTATAGTTCAGC | | | |
| | R2: ATAAACCCTCTTGCAGTTGCATC | | | |
| HEI10$^{mei4}$ | mHei10 atg Forward: ATGTCTTTGTGTGAAGACATGCTGCT | | | No |
| | mei4Genotype Reverse WT: CCCAGCCCCTAGGCACTCAC | WT: 57°C, 35x | WT 300bp | |
| | mei4GenotypeReverse Mut: CCCAGCCCCTAGGCACTCAA | MT: 54°C, 35x | MT 300bp | |
| MLH3$^{tm1Lpkn}$ (null allele) | P1: CGGTTTCCCACCTTCTCTACATCGTCCGTC | 60°C, 35 cycles | WT 250bp, MT 196bp | Yes |
| | P2: TTGGGTAACGCCAGGGTTTTCCCA | | | |

| | | | | |
|---|---|---|---|---|
| | P3: TCAGAGAAGGAAGCCAGTGTCTGCCAC | | | |
| Exo1$^{tm2Wed}$ (null allele) | P1: CTCTTGTCTGGGCTGATATGC | 60°C, 35 cycles | WT 280bp, MT 300bp | Yes |
| | P2: AGGAGTAGAAGTGGCGCGCGAAGG | | | |
| | P3: ATGGCGTGCGTGATGTTGATA | | | |
| Msh2.1$^{tm2.1Rak/J}$ | 12839(184F): TACTGATGCGGGTTGAAGG | 56°C, 35 cycles | WT 211bp, MT 340bp | Yes |
| | 12840(184R): AACCAGAGCCTCAACTAGC | | | |
| | 165R: GGCAAACTCCTCAAATCACG | | | |
| Exo1$^{DA}$ (nuclease dead allele) | Exo1173D InF: CAGGCTGTCATCACAGAGGACTCCGA | 60°C, 35 cycles | WT 611bp, 266bp<br>MT 611bp, 399bp | Yes |
| | Exo1173A InR: ACCTTCTTACAGCCAAATGCGAGGAAGG | | | |
| | Exo1 DA OF: GGACTCTCCTTGCTGACCTTCCATTGTG | | | |
| | Exo1 DA OR: CAGCACCCAAAAAAATCAAACCAAACCA | | | |
| Mus81$^{tm1Chmg}$ | P1: GGTGTGGCCCTGATGGAAGAG | 60°C, 35 cycles | WT 400Bp, MT 370Bp | Yes |
| | P2: GGAGCTAAGGCCTAGCGAGTACAG | | | |
| | P3: CTAGCCGCTTGCGTTCCACAATGT | | | |
| RNF212$^{tm1Nhtr}$ | Common Rev: AACTGTGCATAAGGCCAACC | 60°C, 35 cycles | WT 936bp | No |
| | Forward WT: AGCTCACTGCATAGACCAGGA | | MT 432bp | |
| | Forward MT: CTGTCCATCTGCACGAGACTA | | | |
| FVB-Tg (Stra8-cre)$^{1Reb/LguJ}$ | oIMR7338 Internal control For: CTAGGCCACAGAATTGAAAGATCT | 60°C, 35 cycles | Internal control 324bp<br>Transgene 179bp | Yes |
| | oIMR7339 Internal control Rev: GTAGGTGGAAATTCTAGCATCATCC | | | |
| | oIMR8773 Transgene For: GTGCAAGCTGAACAACAGGA | | | |

| | | | | |
|---|---|---|---|---|
| | oIMR8774 Transgene Rev: AGGGACACAGCATTGGAGTC | | | |
| MLH3$^{D1185N}$ (nuclease dead allele) | For: AAGCCAAGTCTGCATGAGTA | 58ºC, 36 cycles | | N/A |
| | Rev: TAAATGTGCCACTGACTAAAT | | | |
| **Restriction Digest** | Enzyme: Sau96I, at least 4 hours | 37ºC | WT 439bp, 263 bp<br>MT 702 bp | |
| *59.5* PCR | 19HS59.5f16255: GAAAGACGGAAGAGAGCTTCC | 60ºC, 36 cycles | | N/A |
| | 19HS59.5r16825: GGAAGAATAGATGCTTGGTGG | | | |
| **Restriction Digest** | Enzyme: BclI, at least 4 hours | 50ºC | C57BL/6J 377bp, 234bp<br>DBA/2J 616bp | |

**Table S3.** Allele-specific oligonucleotides (ASOs) used for genotyping *59.5*

| ASO | sequence (5'->3') | ASO | sequence (5'->3') | GRCm39 | central polymorphism |
|---|---|---|---|---|---|
| B15102 | AAAAAATTAAAAAAAAAAAAG | Ca15102 | AAAAAATTTTTTTAAAAAAAG | 19:59429036 | rs241448997 |
| B15185 | AAATGTAACGTAGGATAAAA | Ca15185 | AAATGTAATGTAGGATAAAA | 19:59429119 | rs51399465 |
| B15239 | AATGAGGCAGAGGTTGTT | Ca15239 | AATGAGGTAGAGGTTGTT | 19:59429174 | rs48315502 |
| B16003 | TGTACTTTCTGCCTAGTC | Ca16003 | TGTACTTGCTGCCTAGTC | 19:59429938 | rs47029339 |
| B16080 | TACTATCAGTCAATCAATC | Ca16080 | TACTATCAATCAATCAATC | 19:59430014 | rs45707443 |
| B16220 | CCATCAATTCCAAGGAAG | Ca16220 | CCATCAATCCCAAGGAAG | 19:59430154 | rs51260902 |
| B16341 | TCCAATTTCTACCGACTG | Ca16341 | TCCAATTTCTGTCTACCG | 19:59430277 | rs250557647 |
| B16473 | CATATAAACATTTTGCTGT | Ca16473 | CGTATAAATGTTTTGCTGT | 19:59430414 | rs46140223 |
| B16520 | AGTCCAGGCTGGTTTCAA | Ca16520 | AGCCCAGACTGGTTTCAA | 19:59430460 | rs47627339 |
| B16573 | GAACTACCAATCTTCCTG | Ca16573 | GAACTACCAGTCTTCCTG | 19:59430506 | rs51468461 |
| B16582 | TCTTCCTGCCTCTACCTC | Ca16582 | TCTTCCTGTCTCTACCTC | 19:59430516 | rs50736308 |
| B16592 | CTACCTCCTAAATGCTGG | Ca16592 | CTACCTCTTAAATGCTGG | 19:59430527 | rs47233626 |
| B16623 | ACCTCTATGACCAGCTTG | Ca16623 | ACCTCTACGACCAGCTTG | 19:59430558 | rs52052658 |
| B16823 | TCCTGGGCTCCACCAAG | Ca16823 | TCCTGGGAGCCACCAAG | 19:59430758 | rs47310242 |
| B16844 | TATTCTTCCTACTGAGAC | Ca16844 | TATTCTTTCTACTGAGAC | 19:59430779 | rs48231668 |
| B16976 | AAGACATATCTCTCCCAA | Ca16976 | AAGACATGTCTCTCCCAA | 19:59430911 | rs50999333 |
| B17021 | ACGTGTCCCATACTTGAC | Ca17021 | ACGTGTCTCATACTTGAC | 19:59430956 | rs51999729 |
| B17255 | CCACAGATGCAAGCTGCT | Ca17255 | CCACAGAGGCAAGCTGCT | 19:59431190 | rs38558460 |
| B17440 | TGGGACACCAGAAGGTAC | Ca17440 | TGGGACATCAGAAGGTAC | 19:59431375 | rs46966686 |
| B17517 | TCCCACCTATGTCCCCA | Ca17517 | TCCCACCCATGTCCCCA | 19:59431452 | rs51412323 |
| B17558 | AGAAAGTACTCATATGACA | Ca17558 | AGAAAGTGCTCATATGACA | 19:59431493 | rs39149559 |
| B17576 | TGACAGTTTGGCGGTTGG | Ca17576 | TGACAGTTTGGTGGATGG | 19:59431507 | rs49523813 |
| B17583 | TGGACGGATTGGCCAGA | Ca17583 | TGGGCAGATTGGCCAGA | 19:59431522 | rs216052936 |
| B17697 | ATATATGTGTGATGTAGTC | Ca17697 | ATATATGTTTGATGTAGTC | 19:59431631 | rs36601719 |
| B17866 | AATGGCTGAACTGTTGTAG | Ca17866 | AATGGCTAAACTGTTGTAG | 19:59431801 | rs30563970 |
| B17888 | GACCTACTCTAACTCTGG | Ca17888 | GACCTACCCTAACTCTGG | 19:59431823 | rs30719850 |
| B18047 | ATCTCTTTCCCTTTGAGG | Ca18047 | ATCTCTTCCCCTTTGAGG | 19:59431982 | rs48970885 |
| B18142 | AGTGTAGCGGAGCACATC | Ca18142 | AGTGTAGTGGAGCACATC | 19:59432077 | rs51608210 |

| | | | | | |
|---|---|---|---|---|---|
| B18295 | TTATAGGTCTCACTATCCA | Ca18295 | TTATAGGCCTCACTATCCA | 19:59432230 | rs37419451 |
| B18531 | CTTCACATTGACTCTTCCA | Ca18531 | CTTCACATCGACTCTTCCA | 19:59432466 | rs30364053 |
| B18641 | TTACATGTATCTCAGAACT | Ca18641 | TTACATGTGTCTCAGAACT | 19:59432575 | rs31137226 |
| B18723 | ACTGCAGGTGGGGTGGA | Ca18723 | ACTGCAGTTGGGGTGGA | 19:59432658 | rs30661467 |
